# PKA regulates stress granule maturation to allow timely recovery after prolonged starvation

**DOI:** 10.1101/2025.07.06.663161

**Authors:** Sonja Kroschwald, Federico Uliana, Caroline Wilson-Zbinden, Anastasia Timofiiva, Jiangtao Zhou, Sung Sik Lee, Ludovic Gillet, Martina Zanella, Alaa Othman, Raffaele Mezzenga, Matthias Peter

## Abstract

Cells have evolved multiple strategies to survive environmental stress conditions. This includes the formation of membrane-less cytoplasmic ribonucleoprotein structures called stress granules (SGs) that sequester and protect mRNAs encoding many housekeeping genes. SGs are not static biomolecular condensates but transform into solid states during a maturation phase. Although SG maturation is a hallmark of many neurodegenerative pathologies, little is known about the mechanisms and physiological relevance underlying the maturation process. Here we show that yeast SGs mature into a solid-like state during long-term stationary phase stress, which delays SG disassembly and cell cycle restart. Profiling of phosphorylation sites during stationary phase revealed that SG maturation is driven by protein kinase A (PKA)-dependent phosphorylation of the SG proteome. Indeed, upon stationary phase the catalytic PKA subunits condense in SGs where they maintain kinase activity, while during this time cytoplasmic PKA is inhibited. PKA-mediated phosphorylation of key SG components, like the pyruvate kinase Cdc19, is necessary and sufficient for its timely accumulation in SGs, where Cdc19 assembles into amyloid-like structures. Importantly, inhibiting PKA during long-term stationary phase prevents SG maturation, delaying ordered re-start of cell growth after re-feeding. Taken together, these results describe a SG maturation mechanism, selectively activated during chronic stress, that preserves SG integrity and promotes cell survival.

**Graphical Abstract:** 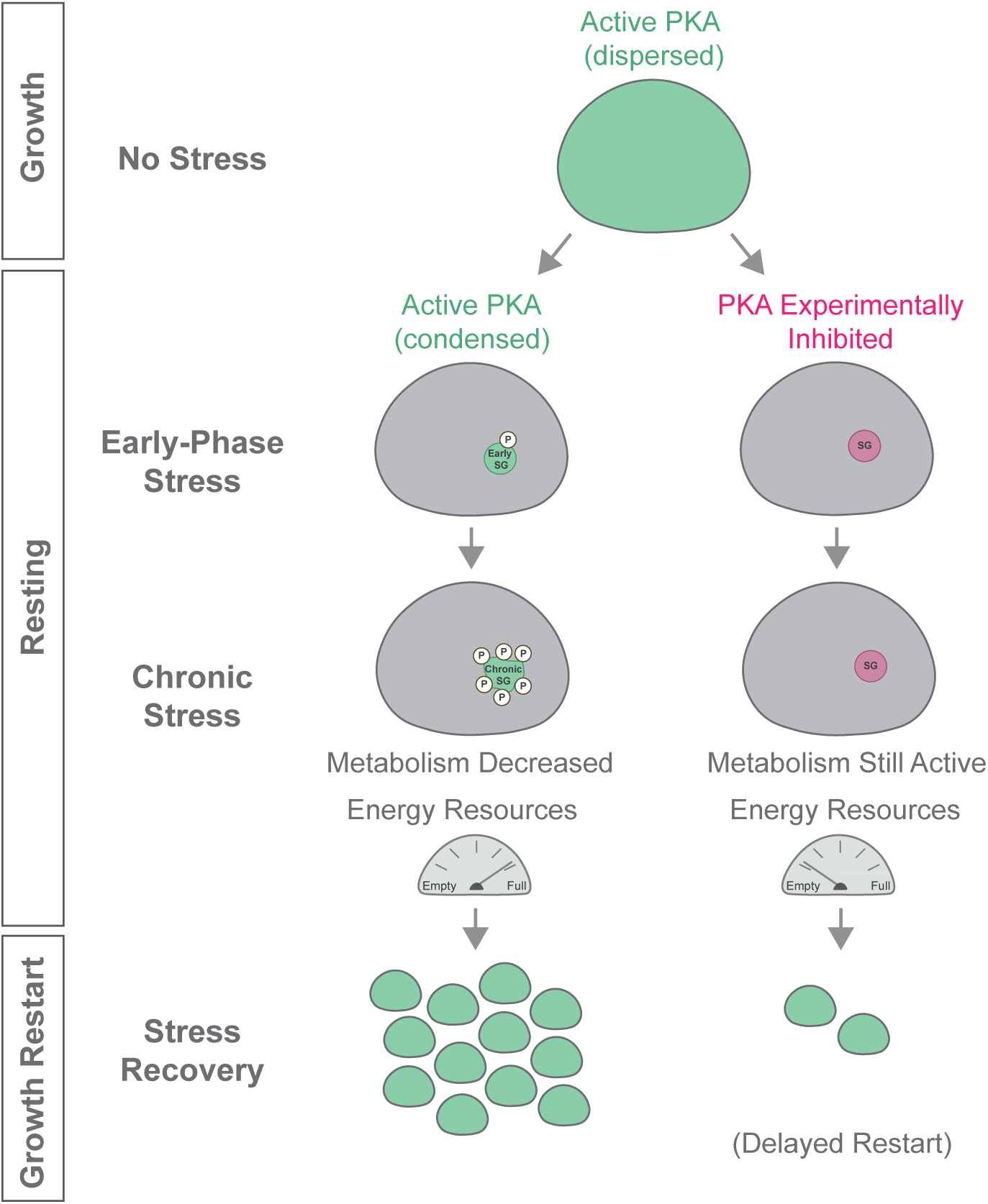

**Highlights:**

- Stress granules mature during chronic stress such as long-term stationary phase
- Phosphorylation of SG components by PKA is a hallmark of SG maturation
- Phosphorylation of Cdc19 by PKA promotes its maturation into an amyloid-like state
- SG maturation acts as a timer for recovery after chronic stress

## Introduction

Cells respond to changing environmental conditions such as nutrient depletion, heat shock and oxidative stress with a variety of evolutionarily conserved strategies. This includes profound rearrangements of their cytoplasm, like viscoadaptation and the assembly of non-membrane bound compartments such as stress granules (SGs) ^1–4^. SGs have been well studied in yeast *S. cerevisiae*, in which acute, transient stress such as e.g. rapid heat shock, provokes an immediate reaction, in contrast to prolonged stress e.g. long-term starvation, which initiates a chronic stress response to prepare cells for long-term cell survival ^5–10^. Studying SGs during chronic stress conditions may be more relevant to unveil their involvement in human diseases that often last months to several years.

With both transient and prolonged stress, SG formation starts by signaling pathways that stall translation initiation, causing ribosomes to run-off their transcripts. Released RNA molecules (mRNAs), associated with RNA-binding proteins (RBPs), such as Pub1 and Ded1, and translation initiation factors then coalesce into SGs that are initially small and dynamic ^3,11–13^. SGs are thought to have a primary function to protect released and sequestered mRNAs and RBPs from degradation, leaving free ribosomes to preferentially engage stress response transcripts to resolve stress induced damage ^11,14,15^. But this simple model of SGs as a transcript reservoir is incomplete. Indeed, despite a global shutdown of translation, only ∼10-15% of bulk mRNA accumulates in SGs and some still undergo translation ^16,17^. Moreover, inhibiting SG formation has no effect on mRNA half-life and cells are still able to inhibit translation of housekeeping proteins ^18–20^. Thus, a more thorough understanding of the physiological functions of SGs will require the dissection of critical components, regulatory mechanisms and cellular phenotypes.

Yeast SGs contain a conserved set of proteins, which may vary with the type of stress and duration ^3,21^. SG components include various metabolic enzymes, such as the pyruvate kinase Cdc19, which assembles into rigid, enzymatically inactive amyloid-like structures ^15,22^. This structural conversion is initiated by a stress-induced decrease in cytosolic pH, which directly regulates protonation of a short amyloid-core motif in Cdc19 ^23^. Importantly, disassembly of Cdc19 aggregates is required to restart ATP synthesis after stress release, thereby coupling cell growth and metabolism ^24^.

In addition to metabolic enzymes, SGs also sequester signaling components such as the Target Of Rapamycin Complex 1 (TORC1) and cyclic AMP-dependent protein kinase A (PKA) ^25,26^. These growth promoting kinases control many cellular processes during exponential growth, including metabolism and cell cycle progression, and their inactivation is important for cells to respond to stress, such as nutrient starvation, and sporulation ^27–32^. PKA is composed of the partially redundant catalytic subunits Tpk1, Tpk2, and Tpk3, which are activated by the Ras-cAMP signaling pathway through dissociating the inhibitory subunit Bcy1 ^29^. During exponential growth the PKA subunits are mostly nuclear and cytoplasmic, while Tpk2, and in part Tpk3, associate with P-bodies and SGs during stationary phase ^26,33^. Thus, SGs may generally sequester and protect essential cellular components to allow stress survival, but it remains obscure why PKA and other signaling components are recruited into SGs.

While SGs are initially small and mobile, they undergo a maturation phase upon prolonged stress conditions, characterized by a liquid-to-solid phase transition, changing their composition, structure and dynamics ^34–36^. As aberrant, matured SGs have been linked to neurodegenerative diseases such as amyotrophic lateral sclerosis (ALS) and frontotemporal lobar dementia (FTLD), it is believed that physiological SGs mature into irreversible, pathological aggregates during disease onset ^34,35,37,38^. However, it remains unclear whether SG maturation itself is merely a feature of an aberrant development, or rather a physiological process to protect SG integrity and function during prolonged stress conditions that can go out of control.

Here, we show that yeast SGs mature into a hardened state during long-term starvation, accompanied by a global increase in phosphorylation of the SG proteome. This conversion is mediated by PKA, which accumulates in SGs and alters their biophysical properties in chronic, long-term starvation, but not acute heat stress conditions. Surprisingly, localized PKA maintains its kinase activity in SGs upon starvation, phosphorylating Cdc19 and other SG components at specific sites, in contrast to cytoplasmic PKA, which is inhibited as expected. PKA-mediated phosphorylation of Cdc19 is necessary and sufficient for timely SG localization and promotes its structural transition into an amyloid-like state. Our data suggest that PKA-mediated SG maturation is functionally important for the ordered restart of cell growth after stress release.

## Results

### Yeast stress granules mature during long-term stationary phase

To study the mechanisms regulating SG maturation, we characterized SG structures marked with mCherry-tagged Ded1 and GFP-tagged Cdc19 during long-term stationary phase stress in *S. cerevisiae*. Starting 24 hours after inoculation during exponential phase, the SG markers Cdc19 and Ded1 began to localize in cytoplasmic foci (Figures 1A and 1B). While the number of SGs stayed constant throughout the time course, their average size increased (Figure S1A), indicating that SGs gradually accumulate macromolecules. Indeed, SG density measured by quantitative phase imaging (QPI)-microscopy increased upon prolonged starvation (Figure 1C). Moreover, fluorescence recovery after photobleaching (FRAP) revealed that the mobility of Cdc19 in SGs decreased during stationary phase (Figures 1D, S1B and 1E), implying SG solidification. Consistent with this notion, sedimentation assays demonstrated that the fraction of soluble Cdc19 decreased with increasing time in stationary phase, in contrast to exponential growth (Figures 1F and quantification 1G). To elucidate whether decreased SG dynamics and solubility are accompanied by structural re-arrangements, we probed the amyloid transition of Cdc19 by measuring its SDS resistance. Indeed, starting at day 5 and 6, but most prominent at day 7 and 8, a SDS-resistant Cdc19 fraction was apparent, characteristic of an amyloid-like state (Figure 1H). As a control, we analyzed the prion Sup35, which showed the expected SDS-resistance in its prion [PSI+] state but not its non-prion [psi-] state at day 8 of stationary phase. Taken together, these results demonstrate that SG foci, marked by Cdc19, change their biophysical properties as they mature in yeast cells exposed to prolonged starvation conditions.

**Figure 1:**
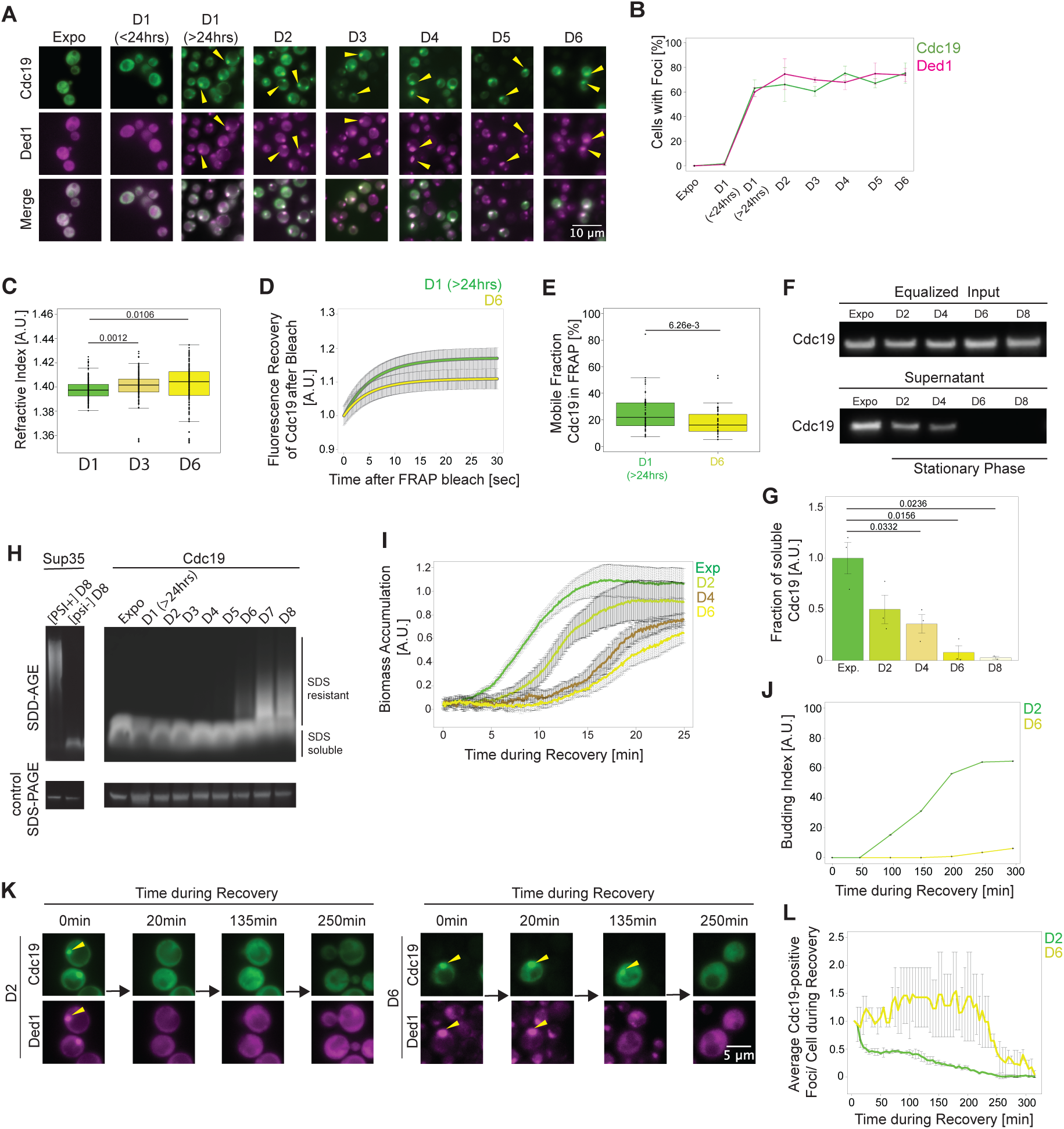
Yeast stress granules mature during long-term stationary phase. **(A)** and **(B)** Fluorescence imaging (**A**) of SGs marked by Cdc19 tagged with GFP (Cdc19) and Ded1 tagged with mCherry (Ded1) and during exponential growth (Expo) and different days in stationary phase (D1-D6). Size bar: 10 μm. The contrast/brightness was adjusted to best visualize cytoplasmic foci. The yellow arrows point to SGs marked by Cdc19-GFP and Ded1-mCherry. Representative images of 3 independent experiments are shown. (**B**) SG formation was quantified by plotting the percentage of cells with Cdc19-GFP and Ded1-mCherry foci. SEM were calculated from at least three independent experiments in 120-1200 cells each condition. **(C)** Refractive index (RI) measurements of SGs visualized by Cdc19-GFP at the indicated days in stationary phase (D1, D3, D6). Statistical significance was determined from 3 independent experiments by an unpaired t-test with equal variance (D1: 244 cells; D3: 177 cells; D6: 187 cells). Statistical relevance determined by an unpaired t-test. (**D**) and **(E)** The dynamics of Cdc19-GFP foci was compared by FRAP analysis in cells after one (D1) or six (D6) days in stationary phase. (**D**) The overlayed curves plot fluorescence recovery (in arbitrary units, A.U.) versus time in seconds (sec) after photobleaching (D1: 36 cells, D6: 31 cells, in 4 replicates each). Error bar shows standard error of the mean. (**E**) The mobile fraction was calculated for each condition, and the statistical relevance determined by an unpaired t-test. **(F)** and (**G**) Extracts prepared from Cdc19-GFP expressing cells during exponential growth (Expo) and different days in stationary phase (D2-D8) were equalized for protein content and then centrifuged. **(F)** An aliquot of the input and supernatant fractions was immunoblotted to visualize Cdc19-GFP. **(G)** The fraction of soluble versus total Cdc19-GFP was quantified for each time point in three independent experiments (D8 only two indep. exp.) and plotted with standard error of the mean. Statistical relevance determined by an unpaired t-test. **(H)** Extracts prepared from Cdc19-GFP expressing cells during exponential growth (Expo) and different days in stationary phase (D1-D8) were probed by SDD-AGE analysis. For control, the prion [PSI+] and non-prion [psi-] states of Sup35 tagged with GFP were analyzed at D8 in stationary phase. SDS-resistant fractions of Cdc19-GFP and Sup35 tagged with GFP were detected by an anti-GFP antibody (n=3 independent experiments). An immunoblot with GFP antibodies was included for control (SDS-PAGE) (n=3 independent experiments). **(I)** Biomass (arbitrary units; A.U.) measured during exponential growth (Exp) and recovery after the indicated days in stationary phase (D2, D4, D6) was plotted against time after refeeding (minutes). Error bars depict standard error of the mean from independent experiments (Exp: n=6; D2: n=4; D4: n=3; D6: n=5). **(J)** The budding index of cells recovering from 2 day (D2) or 6 day (D6) stationary phase arrest was determined microscopically and plotted against time after refeeding in minutes (min) in 100-400 cells each condition. Error bars depict standard error of the mean from 2 independent experiments. **(K)** and **(L)** Cells expressing Ded1-mCherry (Ded1) and Cdc19-GFP (Cdc19) were released from 2 day (D2) or 6 day (D6) stationary phase and **(K)** imaged at the times indicated in minutes (min) after addition of fresh media. Contrast/brightness settings equal between same cell images in same color channel. Size bar: 5 μm. Representative images of 2 independent experiments are shown. **(L)** Foci disassembly was quantified by plotting the average number of foci/cell against time of recovery in minutes (min) in 15-600 cells each condition. Error bars depict SEM from 2 independent experiments.

To examine whether SG maturation has functional consequences, we characterized the recovery of yeast cells that were maintained in stationary phase for different times. Interestingly, after refeeding, the lag time before biomass accumulation increased proportionally with the time spent under starvation conditions (Figure 1I). Although the number of dead cells increased during stationary phase (Figure S1C), the delayed biomass accumulation after refeeding scaled with the re-budding kinetics of individual cells (Figure 1J). Moreover, re-budding correlated with prior dissolution of Cdc19 foci (Figures 1K and quantification 1L), suggesting that cells resolubilize their SGs before cell cycle entry. We conclude that SG maturation during long-term stationary phase leads to slower SG disassembly and delayed cell cycle restart upon nutrient re-addition.

### Stress granule proteins become increasingly phosphorylated during long-term stationary phase

To identify the mechanisms governing SG maturation, we profiled phosphorylation events during long-term starvation. Proliferating cells (Expo) were grown into stationary phase, and samples were harvested at different times from day one to day eight (Figure 2A). As a control, we included cells treated with the TORC1 inhibitor rapamycin, which, like nutrient depletion, triggers cell cycle arrest in G1, but does not lead to the formation of stress granules ^8,15^. After lysis and protein digestion, each sample was split: in one half phosphopeptides were enriched, while the other half was analyzed as full proteome without phosphopeptide enrichment to normalize for protein abundance. In both cases, samples were acquired using a DIA-MS workflow ^39^. We focused on phosphopeptides that exhibited significant intensity level changes (ANOVA, p-value BH-adjusted <0.01 and absolute log2(fold change)>1) compared to the overall mean of all conditions. Following this analysis, we identified 16’908 phosphopeptides belonging to 11’369 phosphosites from 1’564 proteins (∼27% of all yeast proteins, assuming 5’800 total proteins (www.uniprot.org/proteomes/UP000002311)). Among 11’369 phosphosites significantly regulated, 6’595 sites are listed in the BioGRID PTM database (version 4.3.194, ^40^), while 4’774 sites are new and were not previously annotated (Figure 2A). Interestingly, we observed that phosphorylation changes are more pronounced compared to the changes at protein levels, indicating that stationary phase is predominantly regulated by post-translational mechanisms (Figure S2A). Principal component analysis (PCA) revealed a clear separation of samples from exponential growth phase or rapamycin treatment. Moreover, phosphorylation changes also differ among samples taken at different days in stationary phase (Figure S2B), implying a regulated maturation process.

**Figure 2:**
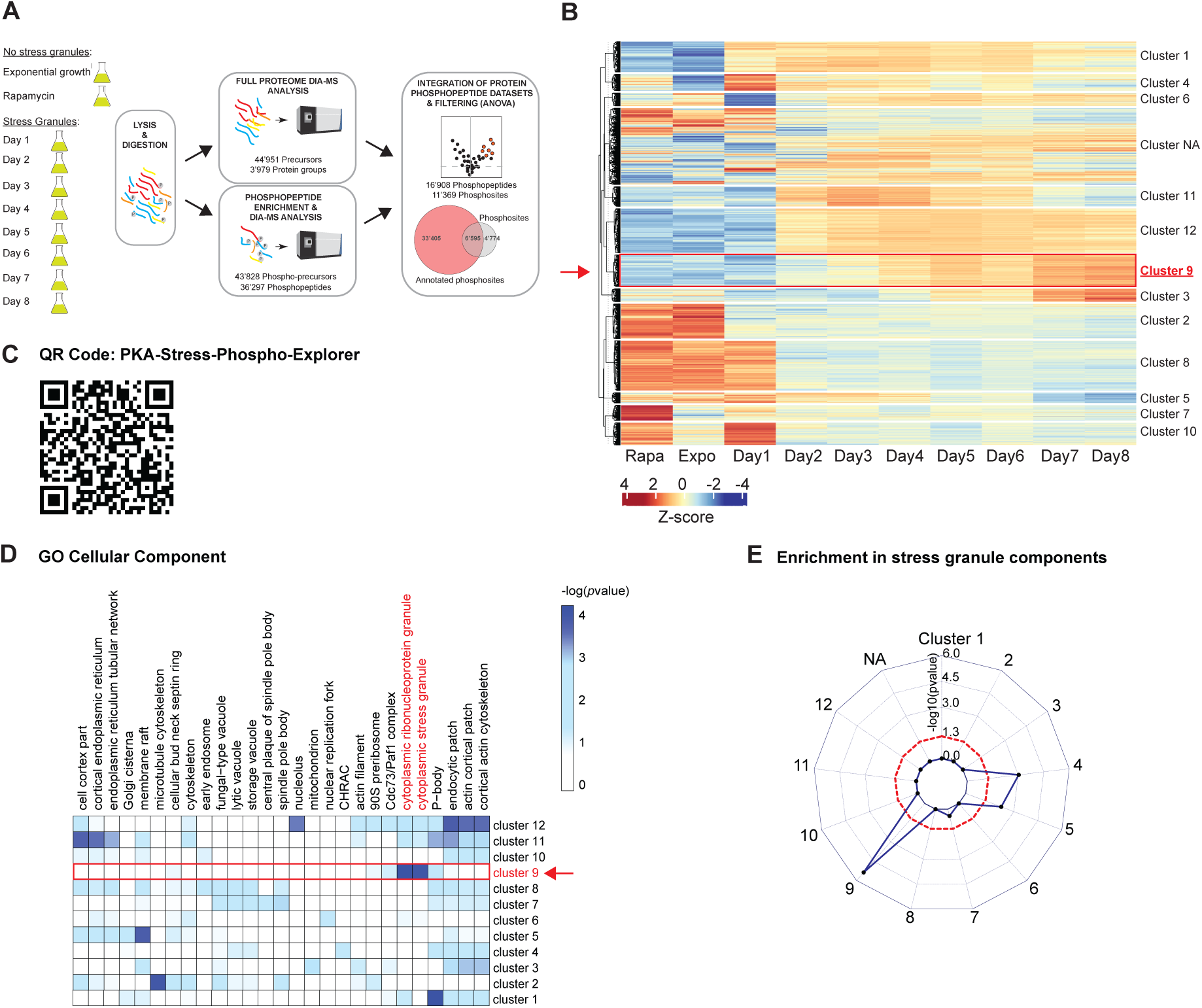
Stress granule proteins become increasingly phosphorylated during long-term stationary phase. **(A)** Workflow of the phospho-enrichment mass spectrometry experiment conducted with cells grown into stationary phase for different days. For control, we also analyzed exponentially growing cells (Expo), and cells arrested with rapamycin (Rapa). Cells were collected in quadruplicates, lysed and proteins digested with trypsin. A fraction of the peptides was directly analyzed, while the remainder was subjected to TiO2 phospho-enrichment. The phosphorylation dataset was filtered for phosphopeptides exhibiting significant changes in one condition compared to the average of all conditions (absolute log2FC>2 and p-value adj<0.05), and then normalized by protein abundance. This analysis identified a total of 16’908 phosphopeptides covering 11’369 phosphosites, 6’595 of which are annotated in the BioGRID PTM database, while 4’774 were previously unknown. **(B)** Heatmap showing normalized phosphopeptide intensities (z-score) for 16’908 filtered phosphopeptides (each phosphopeptide is represented as a horizontal line with the z-score represented by colors (red = increase and blue = decrease over the average abundance of the phosphopeptide)). Unsupervised hierarchical clustering identified 12 clusters (see also Figure S2C). The red arrow and the box highlight phosphosites in cluster 9, which increase during the late stages of the starvation time course. **(C)** QR (Quick response) code to the PKA-stress-phospho-explorer interactive display of the dataset. **(D)** and **(E)** Gene ontology characterization (**D**) of the identified clusters based on the GO Term: Cellular Component. The color intensity represents the (-)log10(*p*value); cluster 9 is highlighted in red. (**E**) Radar plot representing the enrichment of SG proteins in the different clusters based on a SG proteome list generated by BioID proximity labeling analysis ^41^. Statistical significance was tested using a hypergeometric test and is indicated by the red dashed line.

Unsupervised hierarchical clustering based on the profiles of significantly changing phosphopeptides resulted in 12 distinct clusters, ensuring maximum separation between cluster centers (Figures S2C and S2D). The Z-score of each phosphopeptide is displayed in a heatmap (Figure 2B), while Table 1 contains the full dataset listing all phosphoprecursor intensities and phosphopeptide clustering. The dataset is available online (https://pka-stress-phospho-explorer.ethz.ch/, QR code in Figure 2C) with an interactive display to facilitate data visualization. Interestingly, cluster 9 combines 1’238 phosphosites that showed a continuous, gradual increase in intensity over the long-term stationary phase time course. Gene ontology analysis revealed that cluster 9 is enriched in proteins functionally linked to RNA/mRNA-binding (Figure S2E) and characterized as components of cytoplasmic ribonucleoprotein granules and cytoplasmic stress granules (e.g. Cdc19 and Ded1) (Figure 2D), including many identified by proximity dependent ligation experiments (Figure 2E) ^41^. Interestingly, phosphosites in cluster 9 are enriched in intrinsically disordered (IDRs) regions (Figure S2F) ^42^, suggesting that phosphorylation may regulate SG integrity. Taken together, this comprehensive time resolved analysis revealed that many SG components are gradually phosphorylated during stationary phase, thus correlating with SG maturation.

### Protein kinase A (PKA) phosphorylates stress granule proteins during stationary phase

To identify the kinase(s) phosphorylating SG components during stationary phase, we inferred kinase-substrates relationships using the NetworKIN algorithm ^43^. This method predicted potential kinases for all starvation-regulated phosphorylation clusters (Figure 3A), displaying for cluster 9 a significant enrichment for eight kinases (highlighted in red). This analysis was validated via a microscopy screen with fluorescently tagged candidate kinases, which confirmed that from the kinases predicted for cluster 9: Prk1, Tpk2, Tpk3 and Hrr25 assembled cytoplasmic foci during stationary phase, while the other kinases diffusely distributed in the cytoplasm or localized to the nucleus (Figure S3A, highlighted in red). The identification of Tpk2 and Tpk3 is further supported by the presence of the PKA consensus motif (R-R-X-S/T-Y ^44^) in many phosphosites of cluster 9 (Figure S3B), as well as stable or increased Tpk2 and Tpk3 levels during starvation (Figure S3C). While nuclear and diffuse under exponential growth conditions, GFP-tagged Tpk2 and Tpk3 accumulated in SGs during stationary phase (Figures 3B and S3A), in contrast to the inhibitory subunit Bcy1 that remained solely cytoplasmic (Figures 3B and overview PKA life cycle S3D) ^26,33^. Based on these data we speculate that upon recruitment into SGs, PKA progressively phosphorylates SG components during prolonged starvation.

**Figure 3:**
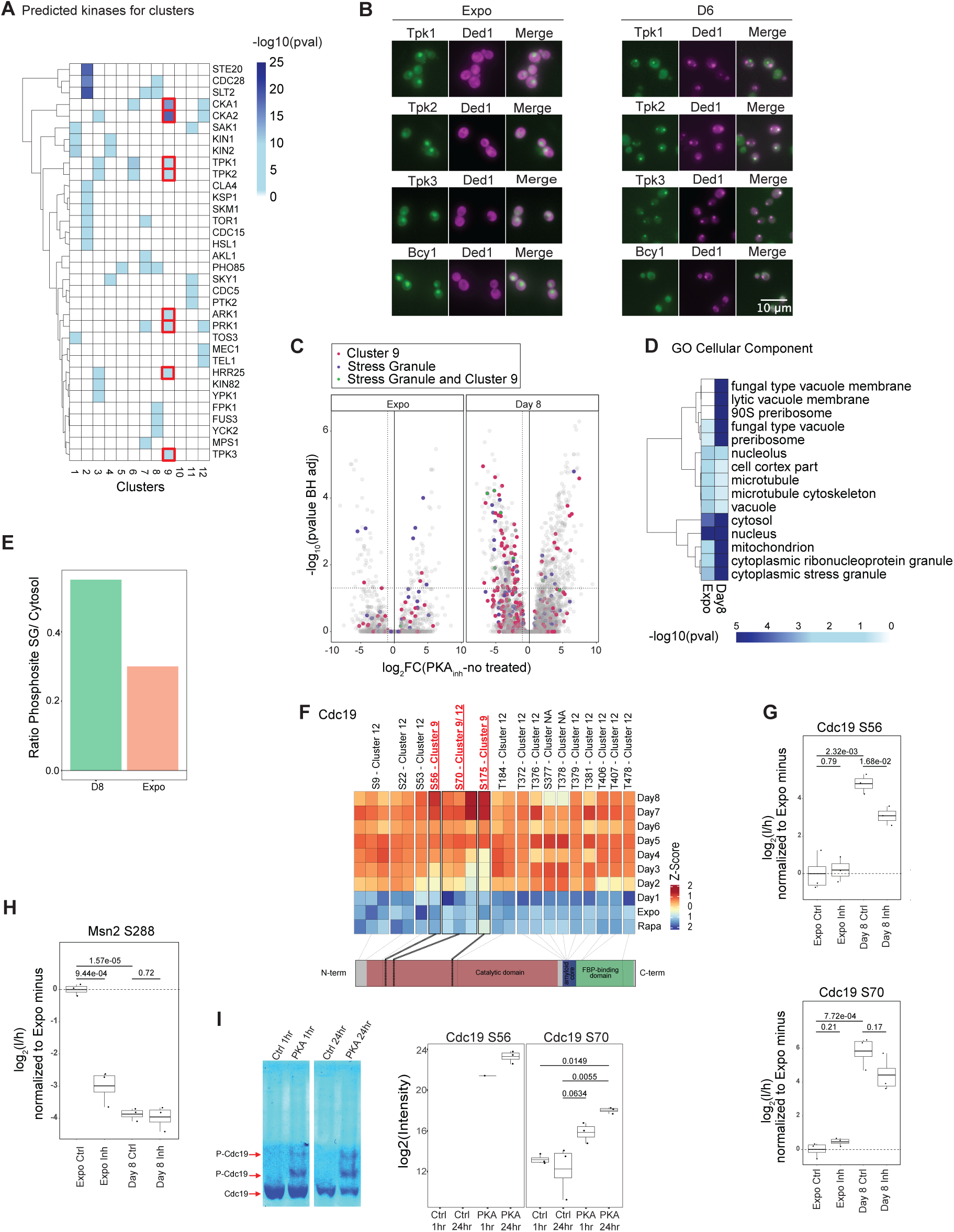
Protein kinase A (PKA) phosphorylates stress granule proteins during stationary phase. **(A)** Substrate-Kinase predictions from the NetworKIN algorithm. The kinases associated with a significant enrichment of phosphosites are displayed for all identified clusters from Figure 2. Candidate kinases significant for cluster 9 are marked in red. **(B)** Fluorescence colocalization analysis of exponential growth (Expo) and stationary phase cells at day 6 (D6) expressing the SG marker Ded1-mCherry (Ded1) and as indicated the catalytic GFP-tagged PKA subunits (Tpk1, Tpk2 or Tpk3) or its regulatory subunit (Bcy1). Note that Tpk2-GFP and in part Tpk3-GFP accumulate in SGs, while Bcy1-GFP remains cytoplasmic. Representaive images of three independent replicates. Size bar: 10 µm. **(C)** Volcano plot showing enrichment and depletion of phosphosites upon treatment with NM-PP1 to inhibit PKA, either in exponential phase (Expo) or after 8 days in stationary phase. Differential abundance is represented on the x-axis, while the significance (p-values adjusted using the Benjamini-Hochberg method) of the changes is shown on the y-axis. Proteins labeled as ’cytoplasmic stress granules’ and/or identified in the experiment in Figure 2A within cluster 9 are highlighted in a different color. Three independent biological experiments were performed. **(D)** Gene ontology characterization of the phosphosites based on the GO term ”Cellular Component” were compared between exponential phase (Expo) and after 8 days in stationary phase (Day 8). The color intensity represents the (-)log10(*p*value). **(E)** Ratio of phosphosites annotated as “cytoplasmic stress granules” and “cytosol” (for this list we excluded proteins annotated as “cytoplasmic stress granules” from the “cytosol” group). The ratio has been calculated in Exponential and stationary phase conditions (D8). **(F)** Heatmap of Cdc19 phosphopeptide intensities (z-score) for the stationary phase time course (day 1 – day 8) and control conditions of exponential growth (Expo) or with rapamycin (Rapa) as originating from experiment in Figure 2A. The annotated cluster of individual phosphosites and their location with respect to functional domains of Cdc19 are indicated. **(G)** Phosphorylation of the cluster 9 Cdc19 sites S56 (upper panel) and S70 (lower panel) were quantified by targeted proteomics in *pka-as* cells during exponential growth (Expo) or after 8 days in stationary phase, with (Inh) or without (Ctrl) addition of NM-PP1. Three independent biological replicates were analyzed. Statistical significance was determined by unpaired t-tests. **(H)** Phosphorylation of the cytoplasmic PKA target site S288 of Msn2 was quantified by targeted proteomics in *pka-as* cells during exponential growth (Expo) or after 8 days in stationary phase, with (Inh) or without (Ctrl) addition of NM-PP1. Three independent biological replicates were considered. Statistical significance was determined by unpaired t-tests. **(I)** Purified Cdc19 was incubated *in vitro* at 30°C with (PKA) or without (Ctrl) recombinant PKA, and phosphorylated Cdc19 was then analyzed on a Phostag-gel stained with Coomassie. Shown are representative kinase reactions of three independent replicates after 1 hr (left gel) or 24 hrs (right gel) incubation time. Red arrows mark Cdc19 species. *In vitro* phosphorylation of Cdc19-S56 (left graph) and Cdc19-S70 (right graph) after 1hr and 24hrs incubation with PKA was quantified by targeted proteomics analysis in three independent biological replicates. Anova significance test was used.

It is generally assumed that PKA is active during exponential phase but inactivated upon nutrient starvation to allow cell cycle exit, and the nuclear import of the transcription factors Msn2 and Msn4 that activate the expression of many stress defense genes ^32,45^. Yet, our results suggest that PKA remains active in SGs during stress conditions. To clarify this apparent paradox, we used a strain harboring an analog-sensitive (as) variant of endogenous PKA (*pka-as*) that can be specifically inhibited by the ATP analog NM-PP1 ^27^. As expected, the bonafide PKA substrate Msn2 accumulated in the nucleus when its phosphorylation was abolished by treating exponentially growing *pka-as* cells with NM-PP1, but rapidly relocalized to the cytoplasm upon washout of the inhibitor (Figure S3E). To directly test whether PKA phosphorylates SG components during starvation but uncouple possible effects of PKA on cell cycle exit, NM-PP1 was added after *pka-as* cells enter starvation (Figure S3F). We confirmed that Msn2-GFP accumulated in the nucleus before NM-PP1 addition, validating that the starved cells had already inactivated cytoplasmic PKA and initiated G1 exit (Figure S3F, upper panel). Moreover, Msn2-GFP remained in the nucleus after prolonged starvation, irrespective of whether PKA was further inhibited by NM-PP1 addition. To explore which phosphosites are regulated by PKA in exponential growing and starved cells, we performed phosphorylation profiling in the presence or absence of the inhibitor NM-PP1. As expected, adding NM-PP1 to exponentially growing *pka-as* cells downregulated predicted PKA phosphosites (Figure S3G). Strikingly, this analysis (Table 2) showed that phosphorylation sites identified in the previous experiment in cluster 9 (Figure 2), and in particular SG components, were phosphorylated in a PKA-dependent manner in stationary phase (8 days), but not during exponential growth (Figures 3C and S3H). Gene ontology enrichment analysis confirmed PKA dependent phosphorylation of cytoplasmic ribonucleoprotein granules, stress granules (GO Cellular Component) (Figure 3D) and components involved in RNA/mRNA binding (GO Molecular Function) (Figure S3I) after 8 days in stationary phase, in contrast to cytosolic proteins (Figure 3E). Together, these results show remarkable PKA dependent phosphorylation of many SG components during stationary phase.

To validate these data, we exploited targeted proteomics (Table 2) to quantify phosphorylation sites in Msn2 and Cdc19. Phospho-enrichment showed that Cdc19 was hyperphosphorylated upon starvation (Figure 3F from dataset presented in Figure 2), showing gradual increase of phosphorylated S56, S70 and S175 in its catalytic domain during stationary phase. Indeed, PRM analysis confirmed that S56 and S70 were phosphorylated during stationary phase and this phosphorylation was dependent on PKA activity (Figure 3G, top and bottom). Surprisingly however, neither S56 nor S70 were phosphorylated by PKA in exponentially growing cells, although cytoplasmic PKA is fully active under these conditions. As expected, PKA-mediated phosphorylation of Msn2 on S288 was high in exponentially growing cells but abolished upon PKA inhibition, whereas Msn2-S288 phosphorylation was low in stationary cells in the presence or absence of NM-PP1 (Figure 3H). Thus, in stationary phase cells PKA activity is maintained in SGs, whereas cytoplasmic PKA is inhibited.

To confirm that phosphorylation of Cdc19 by PKA is direct, we setup an *in vitro* phosphorylation assay using purified components. In contrast to control reactions, additional Cdc19 species could be detected on phos-tag gels after incubating purified Cdc19 with PKA *in vitro* (Figure 3I, left panel), and MS-analysis (Table 3) confirmed phosphorylation of S56 and S70 (Figure 3I, right panel). Taken together, we conclude that during stationary phase, PKA is compartmentalized and remains active in SGs, where it directly phosphorylates SG components, including Cdc19.

### PKA activity is required for stress granule maturation during long-term stationary phase

To test if PKA activity is required for SG maturation, we used *pka-as* strains expressing the SG markers Cdc19-GFP and Ded1-mCherry, and inhibited PKA-activity by NM-PP1 addition after cell cycle exit (as in Figure S3F). Interestingly, accumulation of Cdc19-GFP and Ded1-mCherry in SGs in NM-PP1-treated *pka-as* cells during stationary phase was strongly delayed compared to solvent controls, and this defect was not caused by altered protein levels (Figures 4A and 4B, Figures S4A and S4B). A similar recruitment delay was also observed for the GFP-tagged SG components Pub1, eIF4632, Ksp1, and Rie1 (Figures S4C and S4D), suggesting that PKA-activity promotes SG enrichment of many proteins upon starvation. In contrast, a 10 min or 30 min acute heat stress did not affect SG recruitment of Cdc19 when PKA was inhibited in *pka-as* cells (Figures 4C, 4D and S4E). Indeed, no PKA-dependent phosphosite changes of SG components were detected by MS analysis of heat stressed cells (Figure 4E) ^46^, and no enrichment of phosphosites in proteins of the gene ontology cluster “ribonucleoprotein granules” or “stress granules” was apparent (Figure 4F). Consistent with this observation, Cdc19 was not phosphorylated at S56 and S70 under these acute stress conditions (Figure S4F). We thus conclude that PKA-mediated SG maturation is specific to prolonged stationary phase, but not acute heat stress conditions.

**Figure 4:**
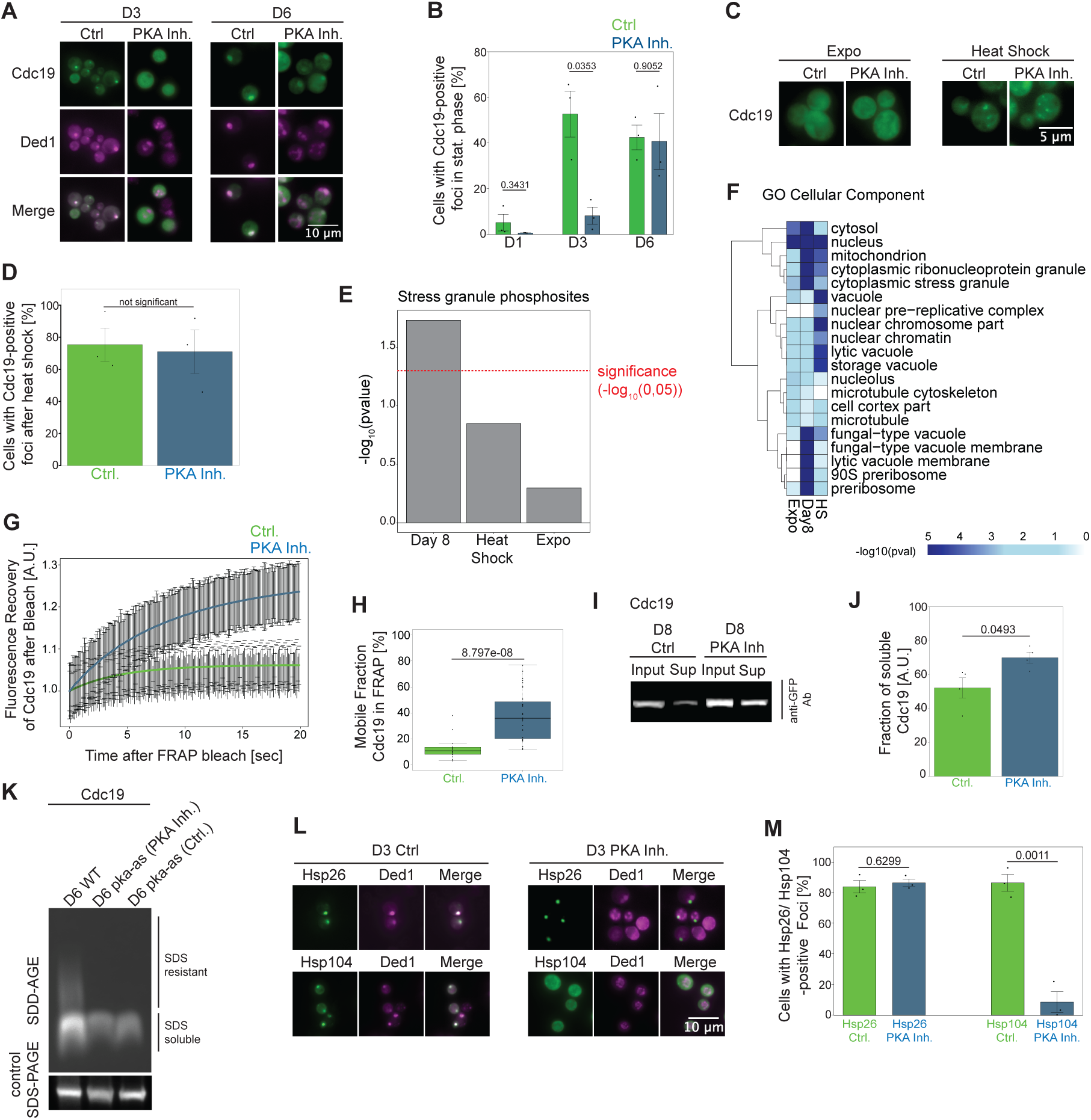
PKA activity is required for stress granule maturation during long-term stationary phase. (A) and **(B)** *pka-as* cells expressing GFP-tagged Cdc19 (Cdc19) and mCherry-tagged Ded1 (Ded1) were analyzed as indicated after 1 day (D1), 3 days (D3) or 6 days (6D) in stationary phase, with (PKA Inh) or without (Ctrl) addition of NM-PP1 to inhibit PKA activity as in the experimental workflow in Figure S3F. (A) Representative microscopy images from 3-4 independent experiments of D3 and D6 stationary phase cells; the contrast/brightness was adjusted to best represent foci. Size bar: 10 µM. (**B**) The percentage (%) of cells with Cdc19-GFP foci were counted at the indicated days in stationary phase. SEM were calculated from three independent experiments analyzing 300-1000 cells for each condition. Statistical relevance determined by an unpaired t-test. (C) and **(D)** Exponentially growing *pka-as* cells expressing GFP-tagged Cdc19 (Cdc19) were heat-stressed (Heat Shock) or not (Expo) for 30 min at 42^0^C, with (PKAInh) or without (Ctrl) addition of NM-PP1 to inhibit PKA activity. (**C**) Representative microscopy images are shown from three independent experiments; the contrast/brightness was adjusted equally between images from the same treatment. Size bar: 5 µM. (**D**) The percentage (%) of cells with Cdc19-GFP foci was counted. SEM were calculated from three independent experiments, analyzing over 1000 cells for each condition. No statistical significance was calculated with an unpaired t-test (p=0.81). (E) Enrichment of phosphosites in proteins annotated in “cytoplasmic stress granules” significantly depleted upon NM-PP1 treatment in D8 condition, Heat Shock (30min, 42C) and Exponential (log2FC< 1 and pvalue BH adj < 0.05). The enrichment was calculated by hypergeometric test considering all proteins identified in the dataset as background, the significance threshold of (- log10(0.05) is marked by the dashed red line. (F) Gene ontology characterization (“Cellular Component”) of the phosphosites significantly depleted (log2FC< 1 and pvalue < 0.05) upon NM-PP1 treatment in Exponential condition, Day8 and Heat Schock condition (30min, 42C). The color intensity represents the (-)log10(*p*value). (G) and (**H**) FRAP analysis quantifying Cdc19-GFP dynamics in 6-day stationary phase *pka-as* cells treated (PKA Inh) or not (Ctrl) with NM-PP1. (**G**) Fitted recovery curves from 16 (Ctrl, green) or 31 (PKA Inh, blue) cells from three independent experiments, shown with standard error of the mean. (**H**) The percentage (%) of mobile Cdc19-GFP was quantified, and statistical significance calculated with an unpaired t-test. (**I**) and (**J**) Extracts prepared from *pka-as* cells expressing Cdc19-GFP after 8 days (D8) in stationary phase and treated (PKA Inh) or not (Ctrl) with NM-PP1 were equalized for protein content and then centrifuged. **(I)** The input and supernatant fraction was immunoblotted to visualize Cdc19-GFP in three replicates. **(J)** The fraction of soluble Cdc19-GFP was quantified. Error bars represent standard error of the mean from four replicates from independent experiments. Statistical significance was calculated with an unpaired t-test (K) SDD-AGE analysis comparing wild-type (WT) and *pka-as* cells expressing Cdc19-GFP starved for 6 days (D6). NM-PP1 (PKA Inh.) or for control DMSO (Ctrl.) was added as indicated. The SDS-resistant fraction of Cdc19-GFP (SDD-AGE, upper panel) and a control immunoblot (SDS-PAGE, lower panel) was probed with an anti-GFP antibody. Note that *pka-as* cells have reduced PKA activity even in the absence of NM-PP1, and thus a comparison to wild-type controls was necessary for this sensitive assay. Three replicates for SDD-AGE and SDS-PAGE were performed in independent experiments. (L) and (**M**) Cells expressing Ded1-mCherry (Ded1) and Hsp26-GFP (Hsp26)) or Hsp104-GFP (Hsp104) were analyzed in 3 days stationary phase (D3) *pka-as* cells treated (Inh) or not (Ctrl) with NM-PP1. (**L**) Colocalization of foci in representative microscopy images of three replicates are shown; the contrast/brightness was adjusted to best represent foci. Size bar: 10µM. **(M)** The percentage (%) of cells with Hsp26-GFP or Hsp104-GFP foci was quantified. SEM were calculated from three independent experiments analyzing 360-440 cells for each condition. Note that SG recruitment of Hsp104 but not Hsp26 requires PKA activity. Statistical relevance determined by an unpaired t-test.

To test whether SGs solidify in a PKA-dependent manner during the maturation process, we performed FRAP analysis of GFP-tagged Cdc19 in starved *pka-as* cells with or without the NM-PP1 inhibitor. Interestingly, Cdc19 mobility in SGs increases upon PKA inhibition (Figures 4G, 4H and S4G), suggesting that PKA inhibits Cdc19 dynamics and promotes SG maturation. Similarly, sedimentation analysis showed that in PKA-inhibited conditions, Cdc19 remained more soluble at day 8 of stationary phase compared to controls (Figures 4I and 4J). Finally, the ability of Cdc19 to assemble SDS-resistant, amyloid-like structures during stationary phase was dependent on PKA activity (Figure 4K). Note that in these assays, the reduced PKA activity in *pka-as* strains was already sufficient to cause the effect even in the absence of the NM-PP1 inhibitor (see “Ctrl” sample).

Since cellular disaggregases recognize and dissolve aggregated structures ^6,12^, we next tested their recruitment to SGs during stationary phase. Interestingly, while the small heat shock protein Hsp26 localized to early SGs irrespective of PKA activity, SG accumulation of the disaggregase Hsp104 was delayed upon PKA inhibition (Figures 4L, 4M and S4H), suggesting that Hsp104 preferentially accumulates in mature SGs. Overall, these results suggest that maturation of Cdc19 and possibly other SG components into a more solid-like state requires PKA activity during stationary phase.

### PKA-mediated SG maturation is important for timely recovery from stationary phase, but not heat stress

To address the functional relevance of PKA-regulated SG maturation we took advantage of the Cdc19-4D mutant, which renders SGs irreversible and as a result starved cells fail to restart growth upon recovery ^15^. We thus set-up a genetic SATAY screen to identify components that allow Cdc19-4D solubilization and enable cells to recover from stress conditions ^47^. Strikingly, many transposons inactivated positive regulators of the PKA pathway (Gpr1, Cyr1, Gpa2) (Figure S5A), including Gpa2, which is known to stimulate the adenyl-cyclase Cyr1 to produce cAMP from ATP ^48^. Indeed, spotting assays confirmed that *cdc19-4D* cells were unable to recover from 6-day starvation stress, while simultaneous PKA inhibition by adding NM-PP1 to *pka-as* strains or *GPA2* deletion suppressed this defect, concomitant with solubilization of Cdc19-4D foci (Figures 5A and S5B). These results suggest that PKA regulates the maturation of SGs during chronic stress, thereby influencing recovery after stress.

**Figure 5:**
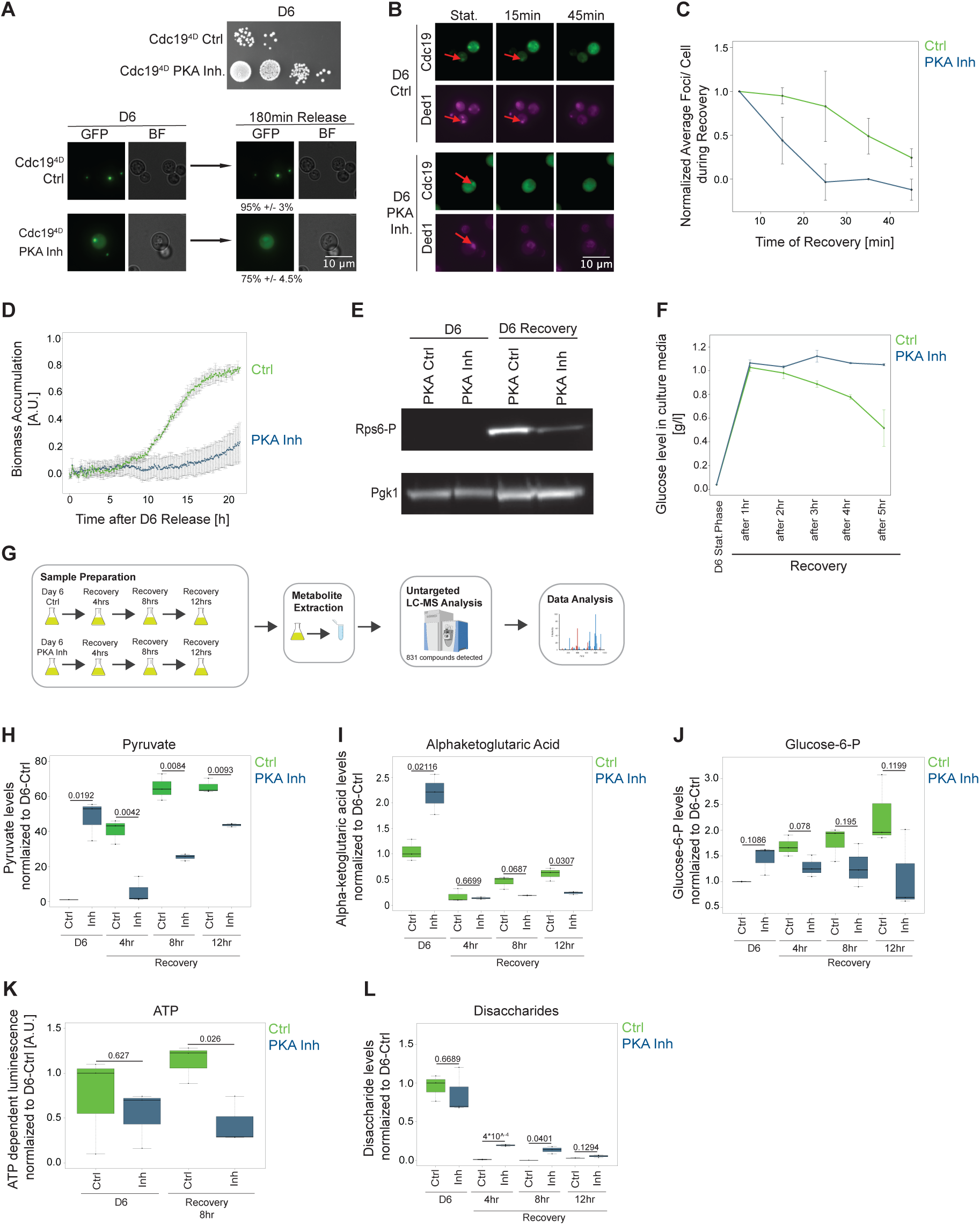
PKA-mediated SG maturation is important for timely recovery from stationary phase, but not heat stress. (A) (Top) Serial dilution spottings on rich media of *pka-as* strains expressing the irreversible Cdc19-GFP mutant (Cdc19^4D^), which were starved for 6 days (D6) and treated (PKA Inh) or not (Ctrl) with NM-PP1 during this time. Representative plate of three replicates is displayed. (Bottom) Bright field (BF) and GFP images (GFP) visualizing Cdc19^4D^-GFP foci in *pka-as* strains during stationary phase (D6) or 180 min after release into fresh medium. Quantification shows percent (%) cells with foci +/- SEM from three independent experiments in 270-300 cells each condition; contrast/brightness was adjusted to best visualize cytoplasmic foci. Size bar: 10 µM. Note that PKA inhibition allows recovery and rapid dissolution of Cdc19^4D^-GFP foci. (B) and (**C**) *pka-as* cells expressing Cdc19-GFP (Cdc19) and Ded1-mCherry (Ded1) were treated (PKA Inh) or not (Ctrl) with NM-PP1 and analyzed microscopically after 6 days (D6) in stationary phase (Stat), and 15 and 45 minutes after releasing the cells into fresh medium without NM-PP1. Images (B) are representative of two independent replicates. The red arrow points to SGs; contrast/ brightness settings are equal between timepoints same condition and same channel. Size bar: 10 µM. **(C)** The average number of foci per cell was quantified at the times indicated (min) after addition of fresh media and normalized to the starting value. Shown with standard error of the mean determined in two independent experiments including at least 50 cells for each condition. (D) Biomass regain (in arbitrary units, A.U.) of *pka-as* cells treated (PKA Inh, blue) or not (Ctrl, green) with NM-PP1 was measured up to 20 hrs (h) into recovery in fresh medium without NM-PP1 after 6 day (6D) stationary phase. Recordings start after inhibitor wash out and addition of fresh media (time 0). Error bars depict standard error of the mean from three independent experiments. Note that inhibiting PKA activity during stationary phase delays recovery. (E) Extracts were prepared from *pka-as* cells treated (PKAas Inh) or not (PKAas Ctrl) with NM-PP1 during 6 days in stationary phase (D6), and after their release in fresh media without NM-PP1 (D6 recovery). TORC1 activity was probed by immunoblotting with an antibody recognizing phosphorylated Rps6 (Rsp6-P). Pgk1 controls equal loading (3 independent experiments). (F) Glucose uptake was determined by measuring glucose levels in the growth media (g/l) of 6-day (6D) stationary phase *pka-as* cells treated (PKA Inh) or not (Ctrl) with NM-PP1, and then analyzed at the indicated hours (hr) in recovery after addition of fresh media without NM-PP1. SEM of two independent experiments are shown. (G) Schematic representation of the metabolomic workflow. Metabolites were extracted from 6-day stationary phase (D6) *pka-as* cells treated (PKA Inh) or not (Ctrl) with NM-PP1 and released into fresh media without NM-PP1 for the indicated recovery time in hours (hrs). Metabolites were extracted and analyzed by untargeted MS. (H) – (**J**) Pyruvate **(H)**, alpha-ketoglutaric acid **(I)**, and glucose-6-phosphate **(J)** levels were quantified by MS analysis in *pka-as* cells at day 6 (D6) in stationary phase with (PKA Inh) or without (Ctrl) NM-PP1, and during recovery for the hours (hr) indicated after addition of fresh media without NM-PP1. The metabolite levels were normalized to D6 Ctrl samples. Statistical significance was determined by unpaired t-test, comparing measurements from three independent replicates. (K) Cellular ATP levels (arbitrary units; A.U.) were measured using a luminescent assay in extracts prepared from 6-day stationary phase (D6) *pka-as* cells treated (PKA Inh) or not (Ctrl) with NM-PP1, and after release for 8 hours by adding fresh media (Recovery 8 hr). The ATP levels were normalized to D6 control samples. Statistical significance was determined by unpaired t-test, comparing measurements from three independent replicates. (L) Disaccharide levels were quantified by MS analysis of 6-day stationary phase (D6) *pka-as* cells with (PKA Inh) or without (Ctrl) NM-PP1, and during recovery for the hours indicated (hr) after addition of fresh media without NM-PP1. Disaccharide levels were normalized to D6 Ctrl samples. Statistical significance was determined by unpaired t-test, comparing measurements from three independent replicates.

To corroborate these results, we next inhibited PKA activity during stationary phase and monitored recovering *pka-as* cells after re-addition of growth media without the NM-PP1 inhibitor. As expected, cells starved for 6 days in the presence of NM-PP1 had more dynamic, less solid-like SGs, which dissolved faster during recovery from stationary phase (Figures 5B and 5C). Surprisingly, however, they exhibited a strong delay in biomass re-accumulation (Figure 5D). In contrast, NM-PP1-treated wild-type cells not carrying the analog-sensitive PKA variant showed no recovery defect (Figure S5C), and exponentially growing *pka-as* cells efficiently resume growth after NM-PP1 washout (Figure S5D), demonstrating that the observed growth restart defect is indeed caused by lack of PKA activity during stationary phase. Interestingly, no delay was observed when cells were stressed by heat shock, confirming distinct mechanisms regulating recovery from acute stress conditions (Figure S5E). Together these results suggest that PKA-dependent SG maturation during stationary phase is required for ordered disassembly and re-activation of cell growth after refeeding.

To investigate the underlying reason for this recovery defect, we analyzed the reactivation of growth pathways and catabolic metabolism. Interestingly, maturation defective cells showed reduced levels of phosphorylated Rps6 (Figure 5E), a 40S ribosomal subunit that serves as a marker for TORC1 activity during growth ^49^. Moreover, PKA inhibition in stationary phase led to reduced glucose consumption during recovery compared to control cells (Figure 5F). To obtain a profile for metabolic flux during the recovery phase, we performed metabolic measurements in *pka-as* cells at three time points (4, 8, 12 hours) after 6 days stationary phase stress in the presence or absence of the PKA inhibitor (Figure 5G, Table 4). The principal component analysis showed a good separation between conditions (Figure S5F). Interestingly, the metabolomic profiles of PKA inhibited cells revealed an accumulation of intracellular pyruvate (Figure 5H), alpha-ketoglutaric acid (Figure 5I) and glucose- 6-phosphate (Figure 5J) at day 6 stationary phase, indicating that glycolysis and the TCA cycle are mis-regulated in the absence of PKA activity during stationary phase (Figure S5G). Upon release into growth supporting conditions, production of these metabolites was delayed in maturation-deficient *pka-as* cells compared to controls, culminating in reduced ATP levels (Figure 5K). In addition, the amounts of accumulated storage disaccharides (e.g. trehalose) (Figure S5G) was comparable at day 6 of stationary phase but decreased slower in PKA-inhibited conditions (Figure 5L). Taken together, these results suggest that PKA-dependent SG maturation is functionally important to control cellular metabolism and allow efficient reactivation of catabolic pathways to produce ATP required for growth after stress.

### Phosphorylation of Cdc19 by PKA during starvation is necessary and sufficient for its SG accumulation and timely recovery from stationary phase

We next examined whether PKA mediated phosphorylation of Cdc19 contributes to SG maturation and the observed phenotypes. To test this hypothesis, we mutated the starvation-induced PKA phosphorylation sites S56, S70 and S175 (Figure 3F) to non-phosphorylatable alanine (Cdc19-3A) and phosphomimicking glutamic acid (Cdc19-3E) residues. As GFP-fusions, both mutants were expressed at similar levels in exponentially growing cells, but Cdc19-3E-GFP seems unstable upon starvation (Figure S6A). Interestingly, Cdc19-3A, but not Cdc19-3E, was able to rescue cell viability in the absence of Pyk2, the paralog of Cdc19 required to substitute essential pyruvate kinase function (Figures S6B and S6C). Neither Cdc19-3A nor Cdc19-3E (with Pyk2 overexpression), produced a dominant effect in exponential growth or upon entry into stationary phase (Figures S6D and S6E), implying that the phosphomimicking Cdc19-3E mutant protein was not toxic. Importantly, however, *cdc19-3A* cells exhibited slower biomass increase during recovery from stationary phase compared to wild-type controls (Figure 6A), while no recovery defect was observed after heat-shock (Figure S6F). Interestingly, *cdc19-3A* cells recovering from stationary phase showed delayed ATP synthesis (Figure 6B), suggesting that PKA-dependent phosphorylation of Cdc19 is functionally relevant for recovery from chronic starvation stress conditions.

**Figure 6:**
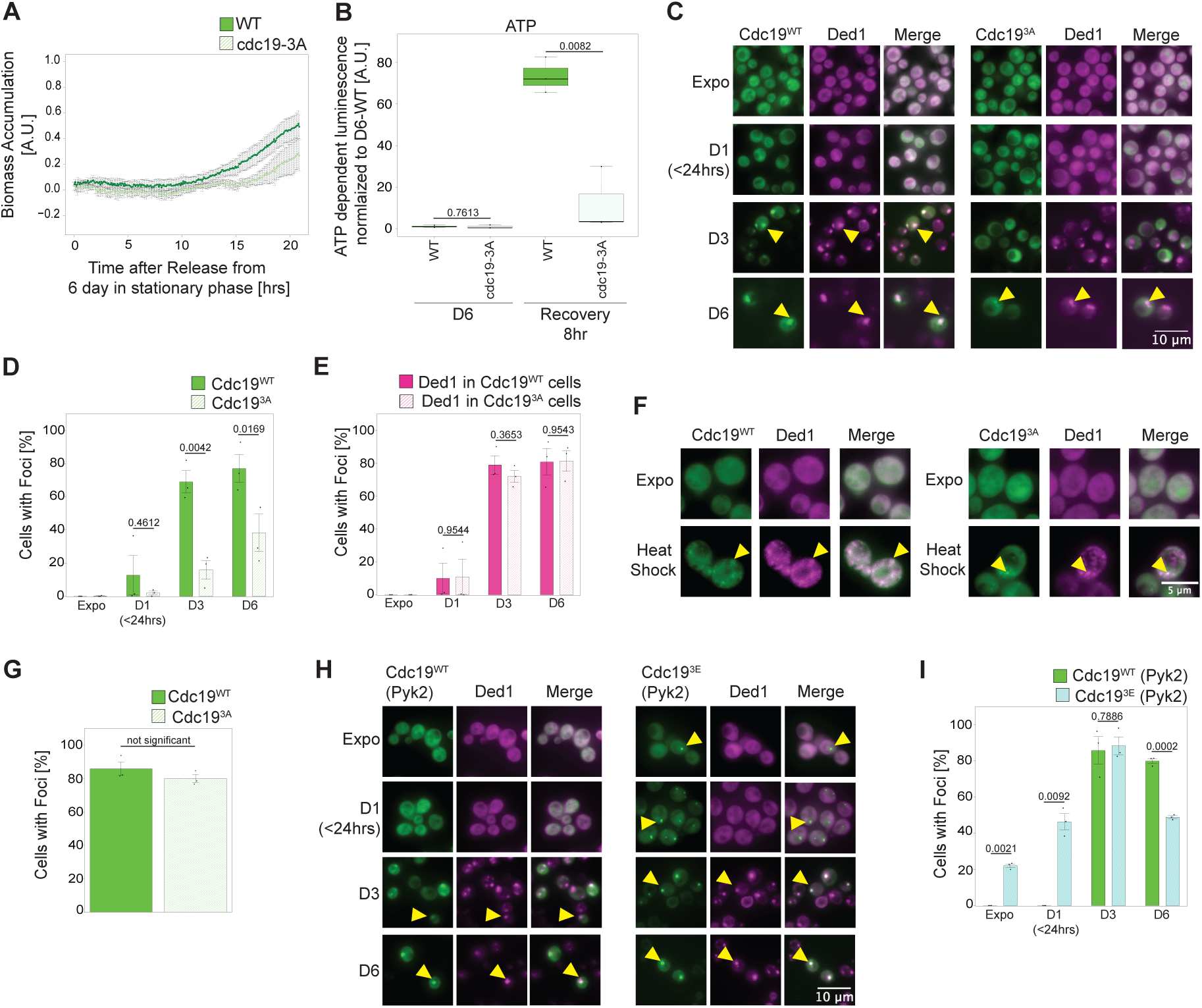
Phosphorylation of Cdc19 by PKA during starvation is necessary and sufficient for its SG accumulation and timely recovery from stationary phase. (A) Biomass accumulation (in arbitrary units, A.U.) of GFP-tagged wild-type (WT) or non-phosphorylatable *cdc19-3A* (S56A, S70A, S175A) mutant cells was measured at different hours (hr) after recovery from 6-day stationary phase. Recordings were started after return to fresh growth media (time 0). Error bars depict standard error of the mean from n=5 independent experiments for WT and n=4 independent experiments for *cdc19-3A* cells. (B) Cellular ATP levels in arbitrary units (A.U.) were compared using a luminescent assay in extracts prepared from wild-type (WT) controls or non-phosphorylatable *cdc19-3A* (S56A, S70A, S175A) mutant cells at 6-day stationary phase (D6) or after release for 8 hours into fresh media (Recovery 8 hr). Statistical significance was determined by an unpaired t-test from three independent replicates. (C), (**D**) and (**E**) Cells expressing Ded1-mCherry (Ded1) and harboring either GFP-tagged wild-type (Cdc19^WT^) or the non-phosphorylatable Cdc19^3A^ (S56A, S70A, S175A) mutant (Cdc19^3A^) were analyzed microscopically when exponentially growing (Expo) or after the indicated days in stationary phase (D1, D3, D6). **(C**) Representative images of three replicates are shown, and the contrast/brightness adjusted to best visualize the cytoplasmic foci at the different days in stationary phase (arrow). (**D**) and **(E)** The percentage (%) of cells with Cdc19^WT^-GFP, Cdc19^3A^-GFP (**D**) or Ded1-mCherry (**E**) (harboring either Cdc19^WT^ or Cdc19^3A^) was quantified, and the standard error of the mean calculated from three independent experiments analyzing 100-900 cells for each condition. Statistical significance was determined by an unpaired t-test. (F) and **(G)** Cells expressing Ded1-mCherry (Ded1) and harboring either GFP-tagged wild-type (Cdc19^WT^) or the non-phosphorylatable Cdc19^3A^ (S56A, S70A, S175A) mutant (Cdc19^3A^) were analyzed microscopically when exponentially growing (Expo) or after heat stress for 30 minutes at 42^0^C (Heat Shock). **(F)** The arrowhead marks cytoplasmic foci. Representative images of three replicates are shown; contrast/brightness adjusted to best visualize the cytoplasmic foci. Scale bar: 5 µM. **(G)** The percentage (%) of cells with Cdc19^WT^-GFP or Cdc19^3A^-GFP foci was quantified, and standard error of the mean calculated from three independent experiments analyzing 170-260 cells for each condition. No statistically significant difference was confirmed by an unpaired t-test (p=0.29). (G) and **(I)** Cells expressing Ded1-mCherry (Ded1) and harboring either GFP-tagged wild-type (Cdc19^WT^) or the phosphomimicking Cdc19^3E^ (S56E, S70E, S175E) mutant (Cdc19^3E^) were analyzed microscopically when exponentially growing (Expo) or after the indicated days in stationary phase (D1, D3, D6). Cell viability was maintained by co-expressing Pyk2. **(H)** Representative images of three replicates are shown; contrast/brightness adjusted to best visualize the cytoplasmic foci at the different days in stationary phase (arrow). **(I)** The percentage (%) of cells with Cdc19^WT^-GFP or Cdc19^3E^-GFP foci was quantified and standard error of the mean calculated from three independent experiments using 300-600 cells for each condition. Statistical significance was determined by an unpaired t-test.

Expression of GFP-tagged wild-type Cdc19 and the Cdc19-3E and Cdc19-3A mutants were used to visualize their localization during exponential growth and upon prolonged starvation. Interestingly, Cdc19-3A-GFP showed delayed SG localization, while the formation of SGs marked by Ded1-mCherry was almost unaffected (Figures 6C-6E). In contrast, no defect in the formation of Cdc19-3A-GFP foci or SGs marked by Ded1-mCherry was observed in heat-stressed cells (Figures 6F and 6G). In contrast, the phosphomimicking Cdc19-3E-GFP mutant analyzed in Pyk2-expressing cells already formed foci during exponential growth and to a higher extent in early stationary phase (Figures 6H, 6I and S6G). No additive defect on foci numbers was observed upon heat stress or in the recovery from heat stress, although the Cdc19-3E containing foci appeared more prominent, most likely because Cdc19-3E is pre-aggregating already in exponential growth (Figures S6H – S6J). Therefore, these results suggest that starvation-induced phosphorylation of Cdc19 by PKA is necessary and sufficient for the timely accumulation of Cdc19 in SGs, with profound implications for recovery from stationary phase.

### Phosphorylation by PKA promotes Cdc19 aggregation

To examine whether phosphorylation of Cdc19 by PKA leads to structural transitions, we first colocalized Cdc19-3E-GFP foci with the mCherry-tagged disaggregase Hsp104 in exponentially growing cells (Figure 7A). In contrast to wild-type controls, Cdc19-3E foci, present in exponential cells, colocalized with Hsp104, implying that Cdc19-3E forms aggregates recognized by Hsp104 even in the absence of stress. Importantly, size exclusion chromatography of cell extracts from exponentially growing cells confirmed that wild-type and Cdc19-3A sediment in lower molecular weight fractions, possibly as monomers and tetramers, while the Cdc19-3E mutant accumulated as an aggregation-prone protein only in higher molecular weight fractions (Figure 7B).

**Figure 7:**
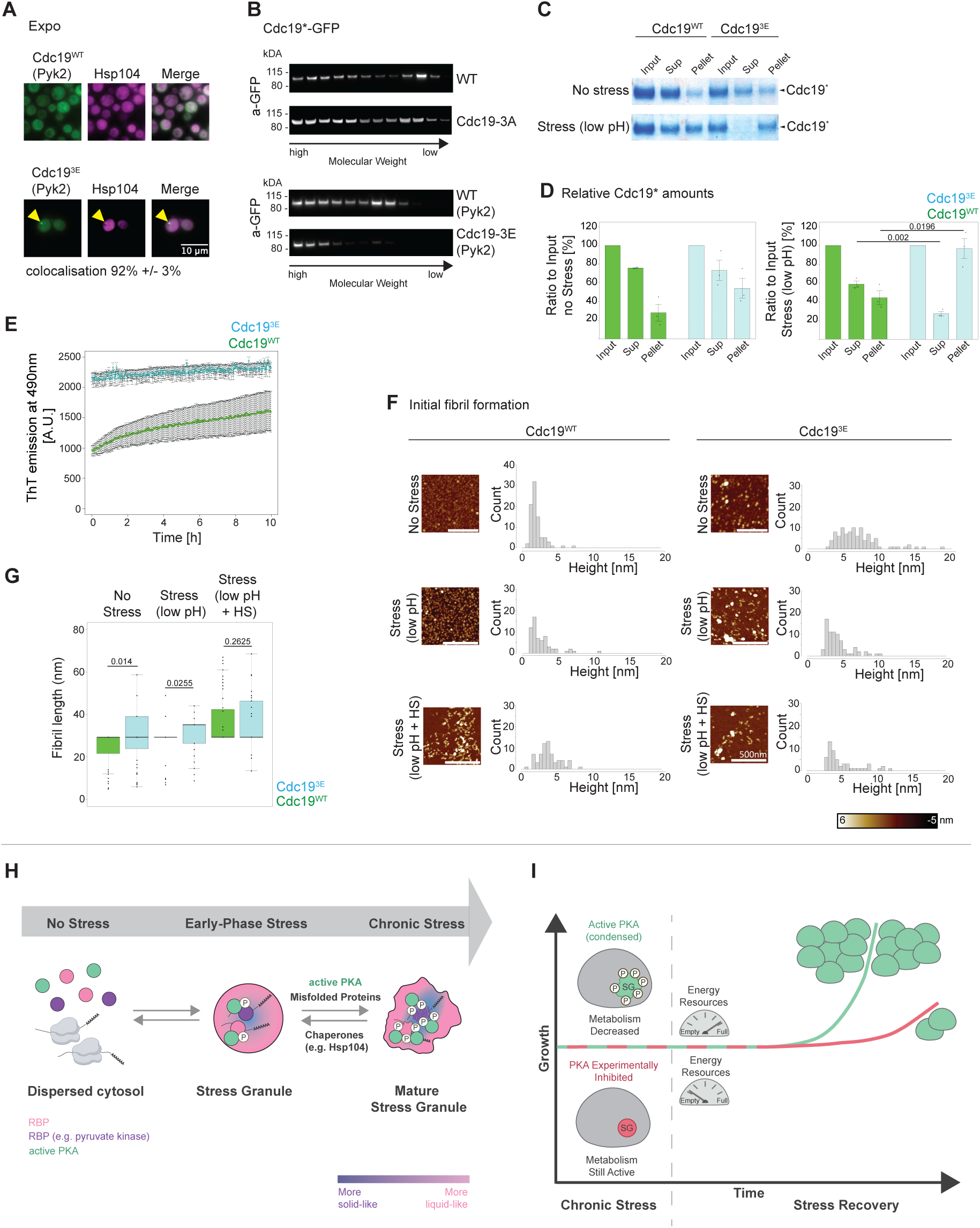
Phosphorylation by PKA promotes Cdc19 aggregation. **(A)** Colocalization analysis of GFP-tagged wild-type (Cdc19^WT^) or the Cdc19^3E^ mutant with the mCherry-tagged chaperone Hsp104 during exponential growth (Expo). Note that the phosphomimicking Cdc193E mutant assembles cytoplasmic foci that in most cases co-localize with Hsp104 (arrowhead). Representative images of three replicates are shown; contrast/ brightness adjusted individually to best represent foci. Numbers represent Cdc19^WT^-GFP or Cdc19^3E^-GFP foci that colocalize with Hsp104-mCherry in more than 200 cells in 4 independent experiments. **(B)** Extracts prepared from exponentially growing cells expressing GFP-tagged wild-type Cdc19 (WT), the non-phosphorylatable Cdc19-3A (S56A, S70A, S175A) or the phosphomimicking Cdc19-3E (S56E, S70E, S175E) mutant proteins were separated by size exclusion chromatography. When indicated (lower panels), Pyk2 was coexpressed (+Pyk2) to maintain viability of *cdc19-3E* cells. The fractions were analyzed by western blotting with an anti-GFP antibody, with larger complexes shown on the left, in two independent experiments for all strains analyzed. Note that Cdc19-3E forms only large assemblies compared to wild-type and Cdc19-3A. **(C)** and **(D)** Recombinant Cdc19^WT^ and Cdc19^3E^ (S56E, S70E, S175E) mutant protein was purified (Input) and analyzed by SDS-PAGE and Coomassie blue staining before and after separation into Supernatant (Sup) and Pellet after no stress or low pH stress exposure (pH=5.9). **(C)** A representative gel image of three replicates is shown. **(D)** Quantification of the protein amount in the Input, Supernatant and Pellet fractions. The relative amounts normalized to the input levels are shown, together with the standard error of the mean from three replicates. Statistical significance was determined by an unpaired t-test. (**E**) Binding of Thioflavin-T (ThT) to recombinant Cdc19^WT^ and phosphomimicking Cdc19^3E^ (S56E, S70E, S175E) mutant proteins was measured by ThT emission at 490 nm (after excitation at 450 nm) at the indicated time points in hours (h), starting after 1hr incubation with TEV protease to cleave the SUMO tag (time 0). Representative curves of three independent replicates with standard error of the mean are shown. **(F) – (G)** Recombinant Cdc19^WT^ and phosphomimicking Cdc19^3E^ (S56E, S70E, S175E) mutant proteins were analyzed by Atomic Force Microscopy (AFM), in unstressed conditions (no stress), after low pH stress (pH=5-6) or strong Stress (low pH + Heat shock (HS)). Shown are representative AFM images and the height (**F**) and length (**G**) profiles of Cdc19 assemblies taken during pre-fibril formation quantified in two independent experiments. Statistical significance was determined by an unpaired t-test. (H) (Model – Part 1) During growth conditions (“No stress”), the catalytic PKA subunits (green) and RNA-binding proteins (RBPs, pink and purple) are dispersed in the cytosol. Ribosomes translate housekeeping mRNAs and stress granules (SG) are absent. Upon nutrient depletion in stationary phase (“Early-Phase Stress”), complexes of RNA and RBPs assemble SGs, and sequester active PKA, while its regulatory subunit Bcy1 remains cytoplasmic. Active PKA in SGs gradually phosphorylate(s) SG components, leading to maturation into a solid state (“Chronic Stress”), together with other factors such as the co-aggregation of misfolding-prone proteins, and counteracted by chaperones such as Hsp104. SG constituents mature into a solid state that is non-dynamic and inactive, and some, like the metabolic enzyme Cdc19, adopt amyloid-like structures. In contrast, the catalytic “active” PKA subunits remain dynamic and active, and thus are in a more liquid-like subdomain. (I) (Model – Part 2) PKA activity inside SGs (top) during chronic stress matures SG components incl. metabolic enzymes, which inhibits catalytic activity and decreases metabolic flux to preserve limited energy resources required to efficiently restart growth after stress subsides. Experimental inhibition of PKA during chronic stress (bottom) prevents SG maturation, enforcing a liquid-like state with residual activity of metabolic enzymes during stationary phase. This increased SG fluidity consumes critical energy resources and as a result delays growth restart.

To test whether increased aggregation of phosphorylated Cdc19 is an intrinsic property, we next compared the sedimentation behavior of recombinant wild-type Cdc19 and the phosphomimicking Cdc19-3E mutant purified as SUMO-fusions from *E. coli* (Figure S7A). The SUMO-tag was then removed by Ulp1-cleavage, and soluble and aggregated Strep-tagged Cdc19 were separated by centrifugation without stress or after a mild pH drop that mimics the physiological stress-induced decrease of cytosolic pH ^2,50,51^. In contrast to wild-type controls, higher amounts of Cdc19-3E were found in the pellet fraction in both conditions (Figures 7C and 7D), suggesting that phosphorylation directly promotes Cdc19 aggregation *in vitro*. Consistent with these results, Cdc19-3E was catalytically inactive (Figure S7B), implying that Cdc19 phosphorylation by PKA promotes its aggregation into catalytically inactive structures.

Kinetic measurements confirmed that purified Cdc19-3E bound high levels of the amyloid-dye thioflavin-T (ThT) already at the onset of the experiment, while wild-type Cdc19 only slowly increased (Figure 7E), suggesting that Cdc19 phosphorylation promotes its amyloid-like conversion *in vitro*. Atomic force microscopy (AFM) was used to compare initial fibril assembly of wild-type and the Cdc19-3E mutant protein *in vitro* (Figures 7F and 7G). Indeed, in contrast to wild-type controls, higher molecular weight Cdc19-3E assemblies were already visible in untreated conditions (“No Stress”). Lowering pH conditions induced the formation of small, uniform globular structures with wild-type Cdc19, while Cdc19-3E assembled larger aggregated species, possibly by promoting lateral interactions of short fibrils. As expected, combining low pH with heat stress induced the formation of small protofilaments of wild-type and Cdc19-3E proteins. Based on this, we conclude that the phosphomimicking Cdc19-3E mutant is prone to adopt an amyloid-like fold and possibly fosters lateral interactions of short fibrils. Thus, Cdc19 phosphorylation by PKA in SGs promotes its maturation into catalytically inactive, amyloid-like structures, which helps cells during recovery from stationary phase.

## Discussion

In this study, we investigated the regulation and function of SG maturation during long-term starvation. We found progressive solidification of SG’s, characterized by specific components such as Cdc19 adopting an amyloid-like state. Surprisingly, many SG components become increasingly phosphorylated by sequestered PKA, revealing an orchestrated physiological process driving SG maturation. Moreover, these findings uncover a new function of PKA in regulating cell survival and recovery from stationary phase but not acute stress conditions.

### Regulation and function of PKA activity in stress granules during starvation

PKA functions during exponential growth, when binding of cAMP dissociates the Bcy1 inhibitor to release active Tpk catalytic subunits. In turn, the catalytic subunits phosphorylate many substrates involved in cell growth and cell division, as well as the general stress transcription factors Msn2 and Msn4, preventing their nuclear localization ^32,48,52^. PKA also inhibits the assembly of P-bodies during exponential phase by phosphorylating the mRNA-decapping factor Pat1, which blocks multimerization of Dhh1 on mRNA ^53^. Upon starvation, decreasing cAMP levels allow Bcy1 binding to the catalytic subunits and PKA inactivation, which triggers cell cycle exit, Msn2/4 nuclear activation and cytoplasmic reorganization ^32,48^. However, while cytoplasmic PKA is inhibited by Bcy1, we found that Tpk2 and Tpk3 retain activity inside SGs during stationary phase and regulate SG maturation by phosphorylating multiple SG components (Figure 7H). This is consistent with findings in starving *S. pombe*, in which deletion of *pka1* impairs SG formation ^54^. Compartmentalized PKA activity in SGs during starvation is demonstrated by SG-specific phosphorylation of’ Cdc19 on S56 and S70. It is striking that these sites are not phosphorylated during exponential growth although cytoplasmic PKA is active and Cdc19 is dispersed in the cytoplasm. Active Cdc19 exists as tetramers, which dissociate into monomers upon nutrient depletion, prone to assembly into amyloid-like fibers ^15^. It is possible that these PKA phosphorylation sites are not accessible in Cdc19 tetramers and are only exposed in Cdc19 monomers or amyloid fibrils. Accumulation in SG’s may simply also increase local PKA concentration, thereby enhancing specific phosphorylation events. This mechanism has been observed in higher eukaryotes, where protein kinase A anchor proteins (AKAPs) enrich PKA holoenzymes in specific subcellular locations ^55^. Finally, PKA may gain additional functionality by binding SG components that extend substrate specificity and/or boost activity ^56^. Thus, controlling the subcellular localization of PKA could be a conserved strategy to control its function in different cellular conditions.

How are Tpk2 and Tpk3, but not Tpk1 and the inhibitory subunit Bcy1, recruited to SGs? Cells lacking all three *TPK* genes are non-viable, but expression of any of the three isoforms rescues viability, suggesting that the catalytic PKA subunits are functionally redundant for cell growth ^57^. However, substrate array analysis revealed striking differences in their substrate preference and specific physiological roles have been reported for each Tpk isoform ^58,59^. Thus, it is possible that Tpk2 and Tpk3 are targeted to SGs by substrate binding. In addition, only Tpk2 contains a glutamine-rich prion-like domain, which is essential for its RNP granule accumulation and efficient RNP granule formation ^60^. Furthermore, recent studies have suggested that the Bcy1-dependent regulation of the catalytic subunits can be isoform specific ^46^. Thus, further work will be required to disentangle isoform-specific functions and spatiotemporal regulation of PKA complexes.

The unexpected discovery that PKA remains active during prolonged starvation challenges the notion that SGs are solely inactive storage compartments with solid-like properties. Like other protein-RNA condensates ^61^, SGs may contain subdomains with different dynamics ^62^ (Figure 7H), providing a cellular environment favorable for ATP-dependent reactions in more liquid-like domains. In addition to PKA, SGs also sequester disaggregating chaperones including Ssa1/Ssa2 and Hsp104 ^6,12^, and together these ATP-dependent processes may shape the biophysical properties and function of this stress-induced compartment.

### Physiological relevance of stress granule maturation

Our results demonstrate that phosphorylation of multiple SG components by PKA regulates SG maturation, characterized by the gradual density increase and immobilization of its constituents. As a result, cells need more time to disassemble SGs and restart growth after stress release, and this lag phase increases with longer starvation periods. Strikingly, PKA phosphorylation of SG components during stationary phase decreases mobile material properties, and individual SG components such as Cdc19 undergo an amyloid-like transition, which is manifested by the formation of SDS-resistant structures. This amyloid-conversion is likely direct, as preventing PKA-mediated Cdc19 phosphorylation leads to increased mobility, while phosphomimicking mutations trigger pre-mature aggregation properties. Moreover, AFM-measurements *in vitro* revealed an increase in lateral pre-fibril clustering, but structural analysis will be required to deduce the underlying mechanism. Available data show that starvation-dependent phosphorylation and dephosphorylation of different sites of Cdc19 regulates multiple assembly states, together with cytosolic pH and allosteric regulators such as FBP ^15,23,24^. Thus, the enzymatic activity of Cdc19 is tightly regulated by spatiotemporal mechanisms including phosphorylation, and its assembly state links cellular metabolism and SG function.

Importantly, SG maturation protects cells during chronic stress conditions, allowing efficient biomass accumulation and ATP production during recovery from prolonged stationary phase. In contrast, SG maturation is not required for cells to survive acute heat stress conditions. Indeed, inhibiting PKA activity during stationary phase delays re-growth upon re-feeding, albeit SGs dissolve earlier. It is possible that untimely release of ATP-consuming proteins from SGs deplete limited energy resources. Similarly, mRNAs may be liberated without re-engaging with translating ribosomes, leading to their degradation. We speculate that orchestrated assembly and dissolution of SGs might help to preserve limited energy resources and critical functions (Figure 7I). Alternatively, SG maturation might contribute to efficiently shut down cellular metabolism under prolonged stress conditions. Indeed, metabolomic analysis revealed that several key metabolites do not decrease as expected during stationary phase when PKA is inhibited (see Figure 6). For example, Cdc19 continues to metabolize PEP to produce pyruvate, which in turn enters the citric acid cycle to produce alfa-ketoglutarate. Continued energy metabolism may thus deplete important resources during starvation, explaining delayed ATP synthesis after glucose re-addition. Thus, PKA may act as a timer to immobilize and structurally organize SG architecture to allow preservation of energy resources during stationary phase (Figure 7I).

### Phosphorylation-driven regulation of stationary phase stress

Although previous work identified condensate and/ or stress related phosphorylation sites, datasets for time-resolved and chronic stress conditions are lacking ^63–66^. Our comprehensive analysis identified over 11’369 phosphosites from 1’564 proteins including 4’774 new and not previously annotated sites, implying that long-term stationary phase is profoundly regulated by post-translational mechanisms. Indeed, PCA not only revealed a clear separation from exponential growth but also distinguishes samples taken during entry and progression through different stages of stationary phase. Most SG components are phosphorylated by PKA, but other kinases might contribute to this complex pattern. Indeed, Hrr25 and Prk1 showed cytoplasmic foci during stationary phase. However, available data shows their accumulation in P-and actin bodies, respectively, rather than SGs ^67^. Thus, future work will be required to disentangle the spatial phosphoregulation of stationary phase, but our time-resolved data set, displayed via the searchable shiny-app, provides an invaluable source to test specific hypotheses.

### SG maturation in mammalian cells: relevant for liquid-to-solid transitions in neurodegenerative diseases?

The finding that yeast SGs mature under chronic stress conditions by phosphorylating many of their constituents prompts the question whether this mechanism could be conserved in mammalian cells. Several studies suggest that mammalian stress granule proteins are controlled by posttranslational modifications ^68–72^. Importantly, accumulating evidence indicates that SGs function as precursors of neurodegenerative disease-associated toxic aggregates ^37,73,74^. Indeed, the disease-associated proteins Fused in sarcoma (FUS) and Tar-DNA binding protein 43 (TDP-43) localize into SGs and their hyperphosphorylation correlates with the development of ALS and FTLD ^68,74–78^. Hyperphosphorylation of proteins in aberrant SGs and disease associated aggregates can affect self-association, interactions with other constituents or cellular localization ^79–81^, suggesting that phosphorylation can increase or decrease aggregation depending on the protein and protein domain targeted ^78–80,82,83^. Thus, a precise understanding of the regulation of SG maturation, and the role of phosphorylation, will be needed. Here, we discovered that phosphorylation of SG components by PKA regulates SG integrity during chronic stress conditions and defects in this physiological mechanism may thus be relevant for disease. Indeed, PKA was implicated in the regulation of the ALS-associated SG protein TAF-15 ^84^. Thus, further studies are warranted to determine the role of PKA for SG maturation in mammalian cells, with possible implications for the liquid-to-solid conversion characteristic for neurodegenerative disorders.

## Data and Code availability

All Proteomics MS raw and analyzed data files, as well as scripts, have been deposited at the Proteomics IDEntifications Database (PRIDE): PRIDE PXD053013 for phosphorylation profile of yeast during starvation; PRIDE PXD052971 for identification of PKA dependent phosphorylation sites and PRIDE PXD052995 for *in vitro* kinase assay of Cdc19 by PKA.

## Author contributions

Conceptualization, S.K. and M.P.; formal analysis, S.K.; funding acquisition, S.K., R.M. and M.P.; yeast analysis and SATAY screening, S.K. and C.W.-Z.; refractive index measurements, S.K. and S.S.L.; proteomics, S.K., F.U., A.T. and L.G.; metabolomics, S.K., M.Z. and A.O.; microscopy: S.K., C.W.-Z., together with ScopeM; protein purification and *in vitro* biochemistry, S.K.; AFM, S.K., J.Z. and R.M.; Visualization, S.K.; Supervision, R.M. and M.P.; writing – original draft, S.K. and M.P.; writing – review and editing, S.K. and M.P. with inputs from Alicia Smith and all authors.

## Acknowledgment

We thank Benoit Kornmann and Agnes Michel for help with the SATAY transposon screen, the ETH Scientific Center for Optical and Electron Microscopy (ScopeM) and in particular Joachim Hehl for assistance with microscopy, and the Functional Genomic Center Zurich (FGCZ) for help with metabolomics measurements. We are grateful to Gabriel Neurohr, Reinhard Dechant, Shady Saad, Gea Cereghetti, Claudia Schmidt and members of the Peter lab for discussions and critical comments on the project. BioRender and Adobe Illustrator were used. This work was supported by the Swiss National Science Foundation (SNSF), the Dementia foundation, ETH Zürich and by postdoctoral fellowships to Sonja Kroschwald from the Human Frontier Science Program (HFSPO) and the European Molecular Biology Organization (EMBO).

## Supplementary Figures and Figure Legends

**Figure S1:**
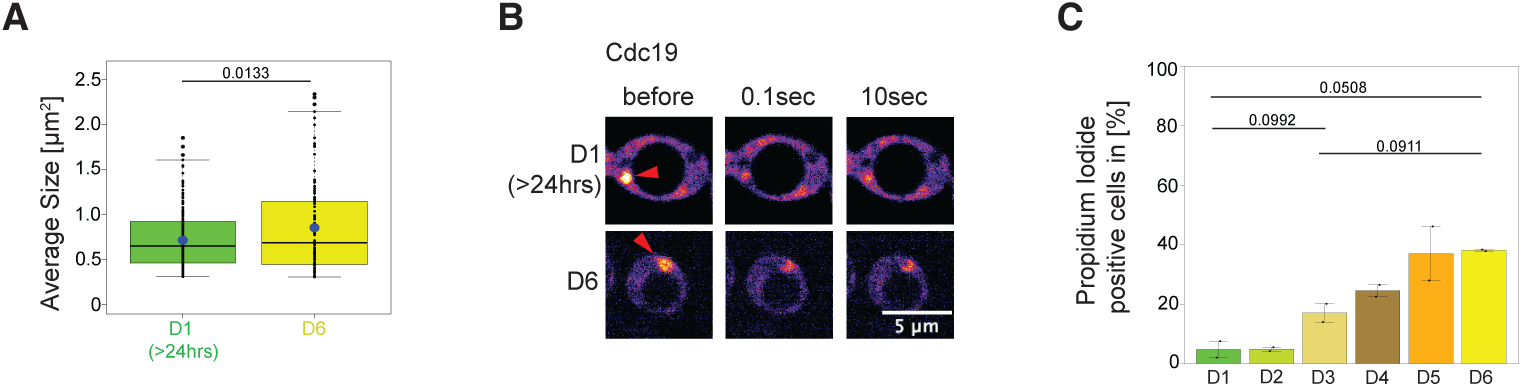
**(A)** Graph depicting average size (μm^2^) of Cdc19-GFP foci at the indicated days in stationary phase (D1: n= 210 foci; 3 replicates, D6: n=105 foci; 5 replicates). Statistical significance was determined by an unpaired t-test. The blue dots mark the means (D1=0.7144 µm^2^, D6=0.8527 µm^2^). **(B)** Representative FRAP images from stationary phase cells on day 1 (D1) and day 6 (D6). The targeted SG marked by Cdc19-GFP (arrow) was imaged before, or 0.1 or 10 seconds after photobleaching. Contrast/brightness settings equal between images for same cell. Recovery of the GFP-signal was quantified in Figures 1D and 1E. Size bar: 5 μm. **(C)** The percentage (%) of dead cells was quantified by propidium iodide staining from up to 6 days of stationary phase (D1-D6). Error bars depict standard error of the mean from 2 independent experiments in 500-1300 cells each condition. Statistical relevance determined by an unpaired t-test.

**Figure S2:**
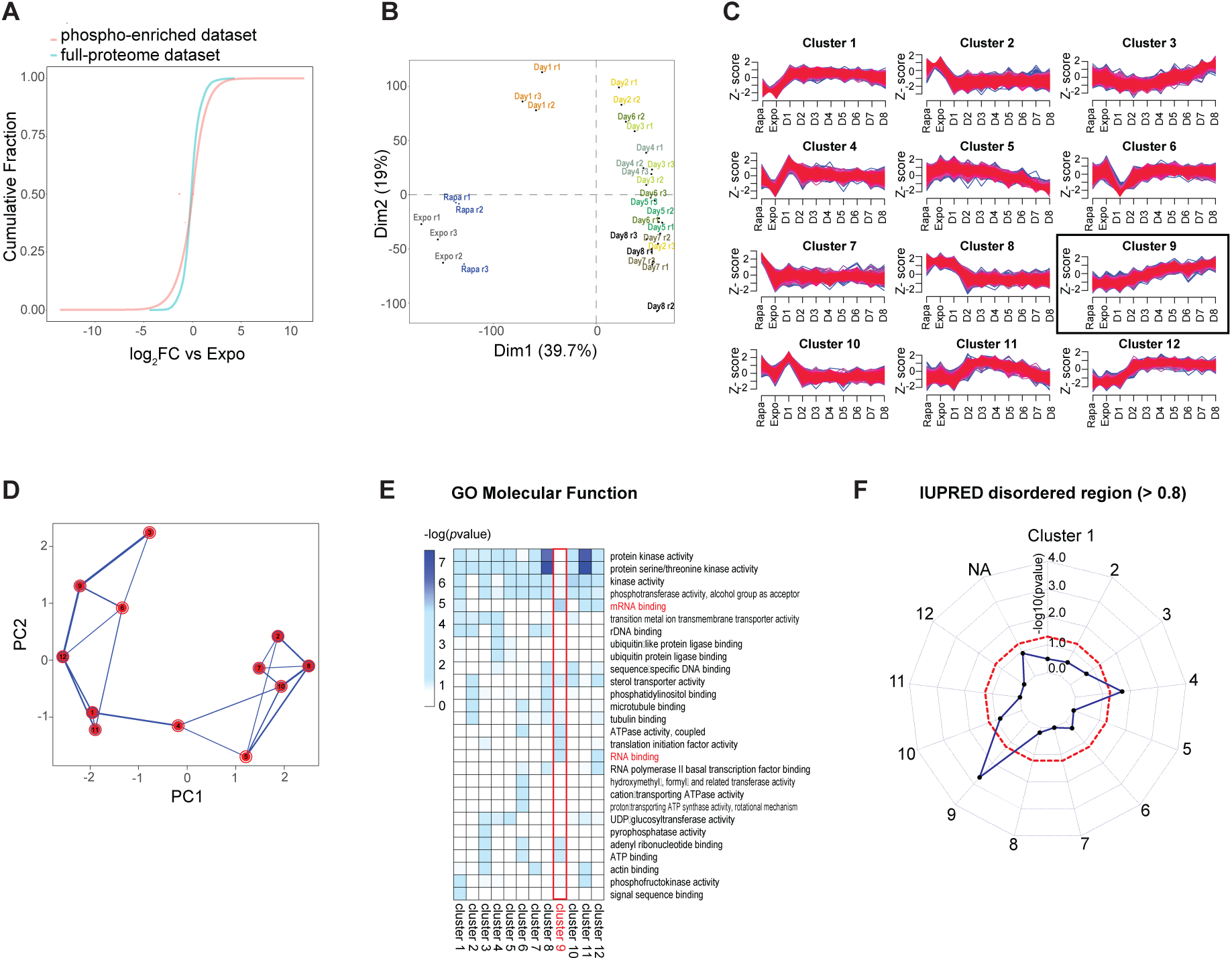
**(A)** Empirical cumulative density function (ECDF plot) for the full-proteome and phospho-enriched datasets derived as described in Figure 2A. The ECDF plot displays the log2 fold change for all conditions compared to exponential growth (Expo), providing a visual representation of their distribution of changes. **(B)** Principal component analysis based on the peptide precursor intensities in the phospho dataset derived as described in Figure 2A. Different conditions are highlighted with distinct colors. **(C)** Plot showing the z-score profile values for clusters in Figure 2B. The stringency of classification is visualized by the line broadness. **(D)** Principal component analysis of the phosphopeptide clusters identified in Figure 2B. **(E)** Gene ontology characterization of the clusters identified in Figure 2B based on the GO Term: Molecular Function. The color intensity represents the (-)log10(*p*value); cluster 9 and RNA associated terms are highlighted in red. **(F)** Radar plot testing the enrichment of phosphorylation sites in disordered regions in the clusters identified in Figure 2B. Intrinsically disordered regions are defined as protein region comprising at least five amino acids with an IUPRED score higher than 0.8. Statistical significance was assessed using a hypergeometric test and indicated by the dashed red line.

**Figure S3:**
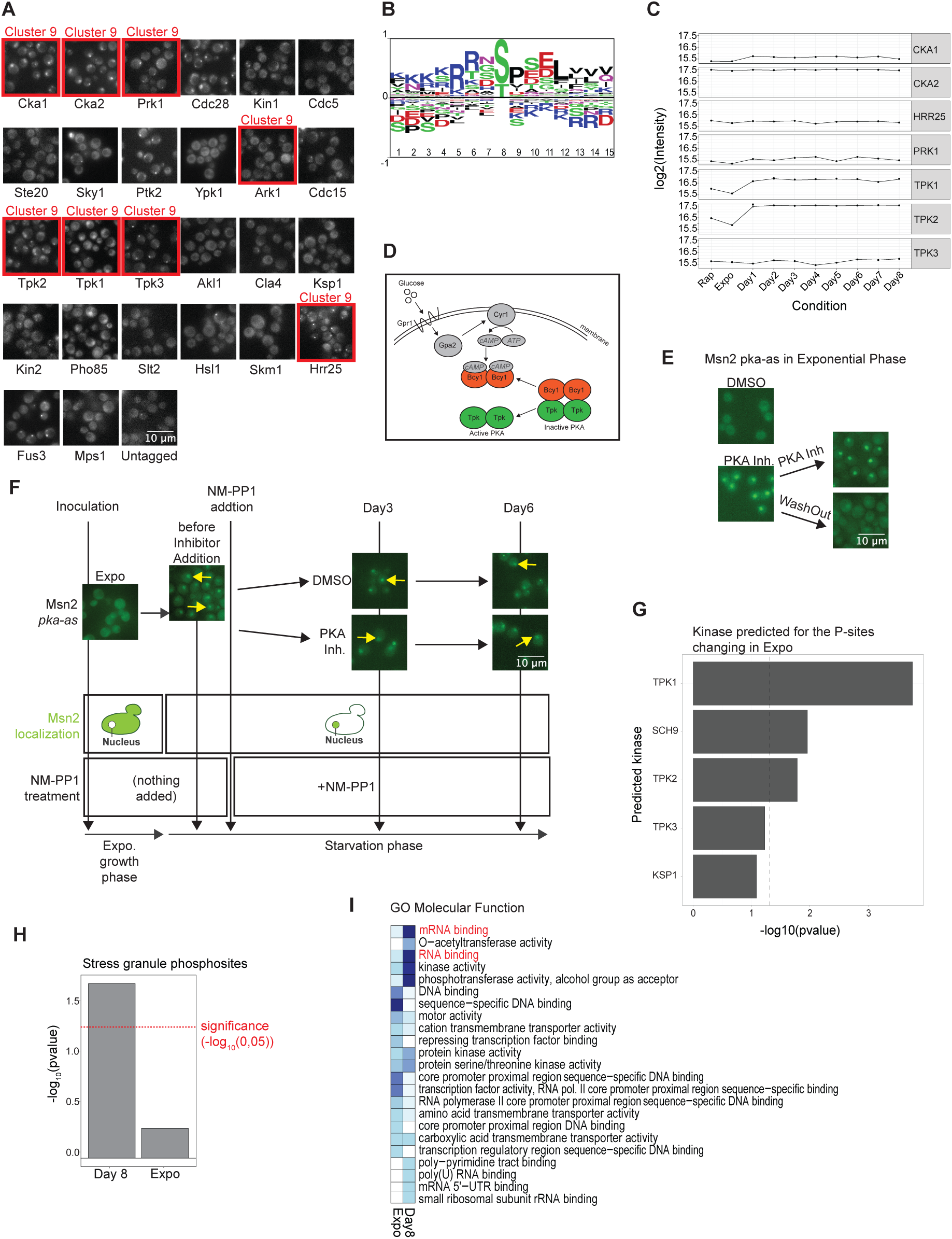
**(A)** The localization of the indicated GFP-tagged protein kinases was examined in a fluorescence microscopy screen in cells arrested in stationary phase for 1 day. Predicted kinases (Figure 3A) for cluster 9 phosphosites (Figure 2) are marked in red. Representative images of 3 independent experiments. **(B)** A consensus motif encompassing all phosphorylated serine and threonine residues grouped in cluster 9 from Figure 2 was determined. Note that the motif resembles the substrate consensus motif of PKA, which is “R-R-X-S/T-X” ^44^. **(C)** Protein levels (log2(intensity)) for the predicted kinases shown in Figure 3A were quantified by MS-analysis at different days in stationary phase (Day 1-8), and control conditions of exponential growth (Expo) or with rapamycin (Rapa). Peptides for Ark1 were not detected. **(D)** Schematic illustration of the yeast PKA pathway with upstream regulators. PKA is active in high glucose/nutrient conditions when Gpa2 activates Cyr1, converting ATP into cAMP, which in turn triggers dissociation of Bcy1 from the catalytic Tpk subunits. **(E)** The localization of GFP-tagged Msn2 was analyzed microscopically in exponentially growing *pka-as* cells. The cells were treated for 30 min with NM-PP1 to inhibit PKA (PKA Inh) or its solvent DMSO for control. NM-PP1 was then washed out (washout) or not (PKA Inh) and imaged again 30 min later. Contrast/brightness settings equal between images before and after washout. Size bar: 10 µM. Representative images of three replicates, respectively are shown. Note that PKA inhibition triggers nuclear accumulation of Msn2-GFP, which is readily reverted after NM-PP1 washout allowing PKA re-activation. **(F)** Workflow to examine PKA-dependent maturation processes during starvation. Exponentially growing *pka-as* cells expressing Msn2-GFP were allowed to enter stationary phase, at which time the NM-PP1 inhibitor (PKA Inh) or for control DMSO (DMSO) was added for 3 or 6 days in stationary phase. Msn2-GFP was analyzed microscopically before and after addition of NM-PP1, and its localization reporting on PKA activity is shown in the GFP-images and schematically indicated below. Contrast/brightness settings equal between images from same timepoint. Note that, as expected, Msn2-GFP is already nuclear by the time of NM-PP1 addition due to nutrient starvation and remains nuclear throughout stationary phase (arrows), irrespective of the presence or absence of the PKA inhibitor. Representative images of three replicates for “Expo”, “before Inhibitor Addition”, Day3 (DMSO and PKA Inh.) and Day6 are shown. **(G)** Substrate-Kinase predictions from the NetworKIN algorithm in exponential condition (Expo). The kinases associated with a significant enrichment of phosphosites that are depleted in exponential condition upon NM-PP1 inhibitor treatment are shown. **(H)** Enrichment of phosphosites in proteins annotated in “cytoplasmic stress granules” significantly depleted upon NM-PP1 treatment in Exponential and D8 condition (log2FC< 1 and pvalue BH adj < 0.05). The enrichment was calculated by hypergeometric test considering all proteins identified in the dataset as background, the significance threshold of (-log10(0.05) is marked by the dashed red line. **(I)** Gene ontology characterization (“Molecular Function”) of the phosphosites significantly depleted (log2FC< 1 and pvalue < 0.05) upon NM-PP1 treatment in Day8 and Exponential condition. The color intensity represents the (-)log10(*p*value).

**Figure S4:**
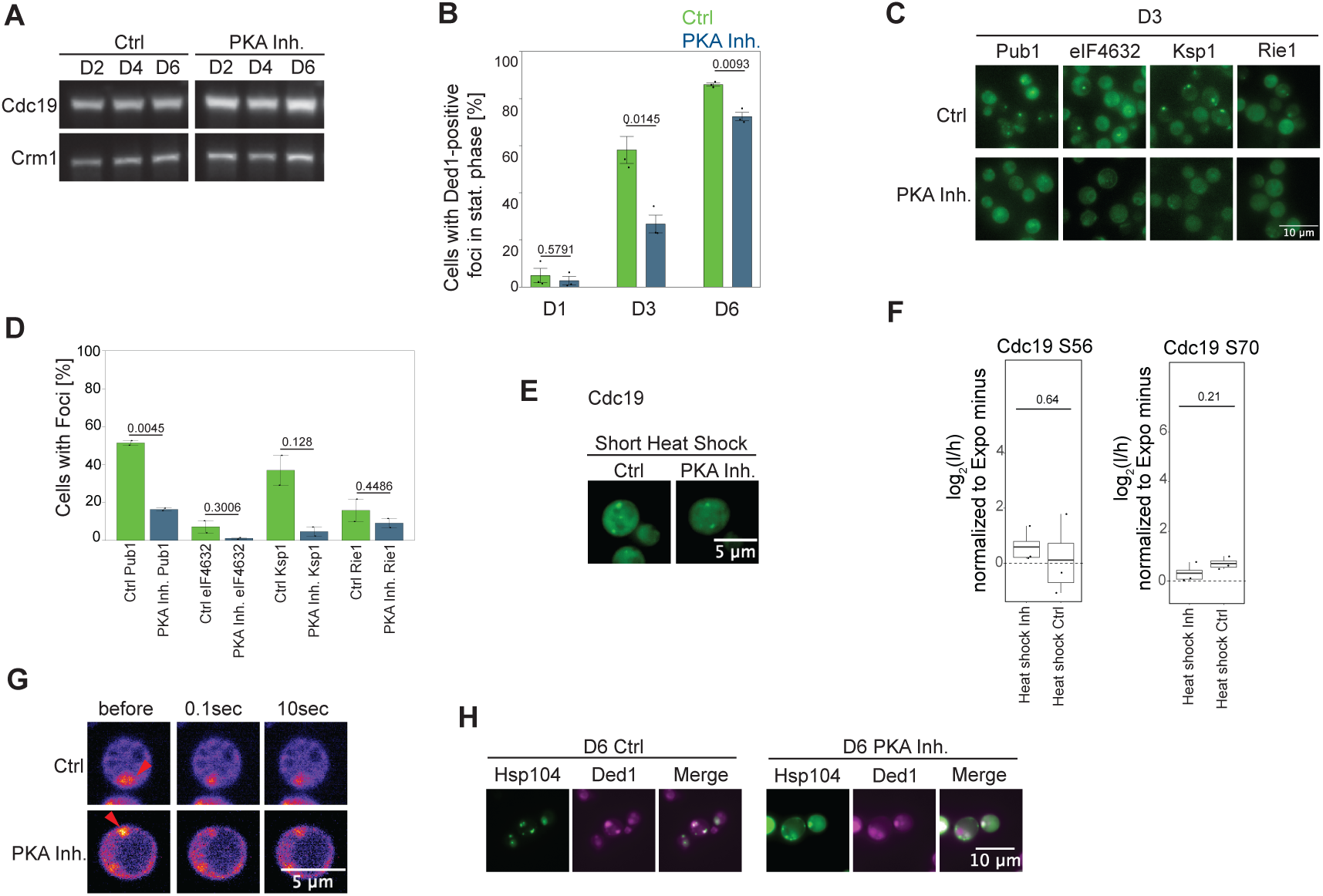
**(A)** Cdc19-GFP expression was compared by immunoblotting of cell extracts prepared from *pka-as* cells treated (PKA Inh) or not (Ctrl) with NM-PP1 after 2 (D2), 4 (D4) or 6 (D6) days in stationary phase. An antibody recognizing Crm1 controls equal loading. Two replicates from independent experiments. **(B)** Quantification of the percentage (%) of Ded1-mCherry positive foci in *pka-as* cells treated (PKA Inh) or not (Ctrl) with NM-PP1 after the indicated days in stationary phase (D1, D3, D6). SEM were calculated from three independent experiments in 400-900 cells each condition. Statistical relevance determined by an unpaired t-test. **(C)** and **(D)** The localization of GFP-tagged SG components (Pub1, eIF4632, Ksp1 and Rie1) in *pka-as* cells treated (PKA Inh) or not (Ctrl) with NM-PP1 were analyzed after 3 days (3D) in stationary phase. (**C**) Microscopy visualizing GFP-tagged Pub1, eIF4632, Ksp1 and Rie1; equal contrast/brightness between images visualizing same protein. Size bar: 10µM. **(D)** The percentage (%) of cells with foci was quantified from two independent replicates in 300-800 cells each condition and is shown with the standard error of the mean. Statistical relevance determined by an unpaired t-test. **(E)** Cdc19-GFP foci were imaged in *pka-as* cells treated (PKA Inh) or not (Ctrl) with NM-PP1 and exposed to a short heat-stress (10 min, 42^0^C). Representative images of two replicates, equal contrast/brightness settings apply. Note that a smaller area was cropped to better highlight heat shock foci. Size bar: 5µM. **(F)** Phosphorylation of the cluster 9 Cdc19 sites S56 and S70 quantified by targeted proteomics in *pka-as* cells treated (Inh) or not (Ctrl) with NM-PP1 and exposed to heat stress for 30 min at 42^0^C. Statistical significance assessed by unpaired t-tests. **(G)** Representative FRAP images of Cdc19-GFP foci in 6 days stationary phase *pka-as* cells treated (PKA Inh) or not (Ctrl) with NM-PP1. Cdc19-GFP localizing into foci (arrow) was imaged before, or 0.1 or 10 seconds after photobleaching. Recovery of the GFP-signal was quantified in Figures 4G and 4H. Representative images of three independent experiments are shown; contrast/ brightness settings are equal between images same condition. Size bar: 5 µM. **(H)** Colocalization of Ded1-mCherry (Ded1) and Hsp104-GFP (Hsp104) was analyzed in 6 days (D6) stationary phase *pka-as* cells treated (PKA Inh) or not (Ctrl) with NM-PP1. Representative microscopy images of three replicates are shown; contrast/brightness was adjusted to best visualize cytoplasmic foci. Scale bar: 10µM

**Figure S5:**
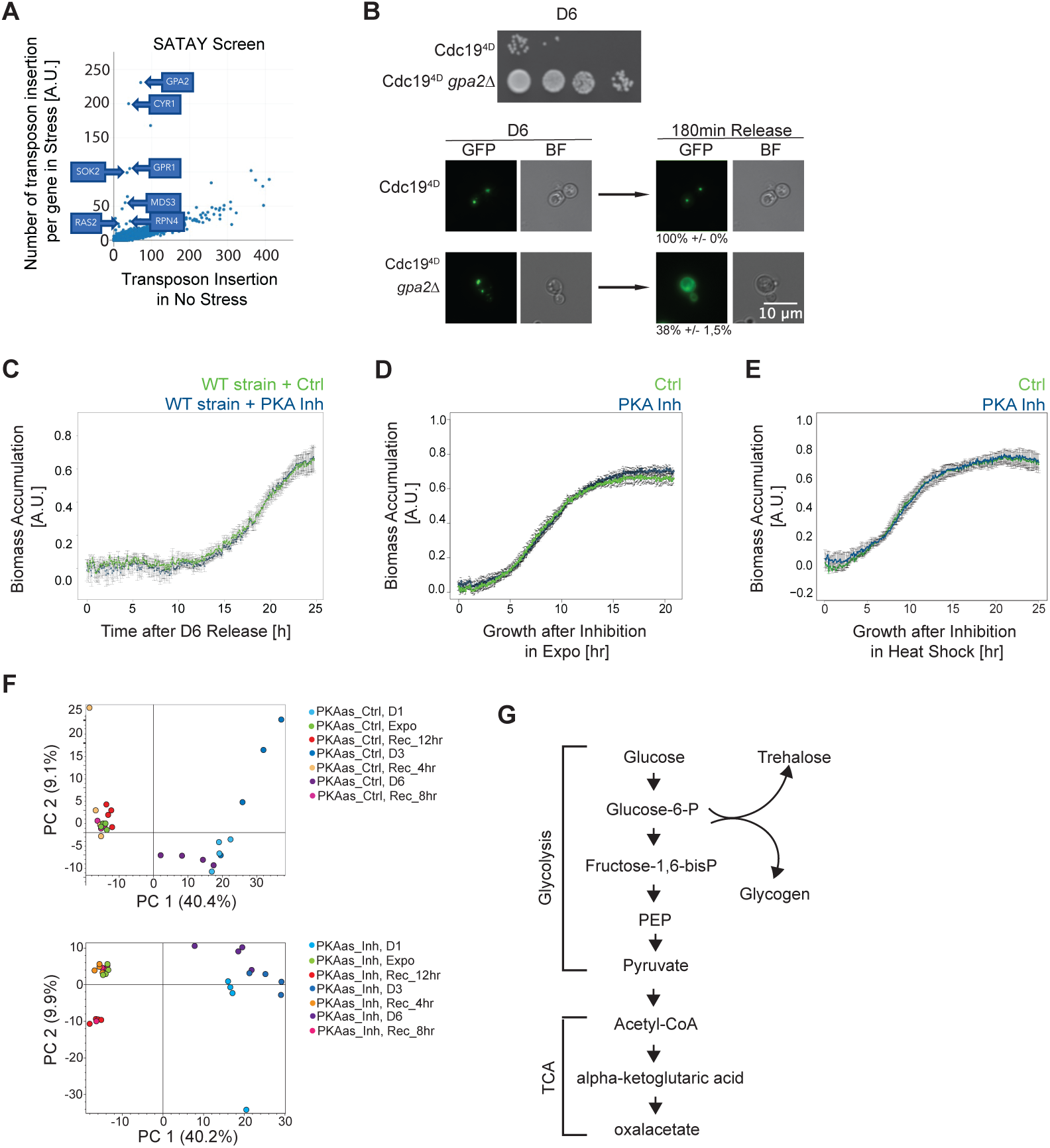
**(A)** A SATAY transposon screen identified genes that upon deletion through transposon insertion rescued cells expressing the irreversible Cdc19-4D mutant after heat stress (HS). Eight million clones from control versus HS sample were analyzed. Note that most identified genes are known regulators of the PKA pathway (boxes). **(B)** (Top) Cdc19^4D^-GFP expressing cells deleted (*gpa2τ<*) or not for *GPA2* were starved for 6 days (D6) and then spotted in serial dilution on nutrient-rich media. (Bottom) Bright field (BF) and GFP images (GFP) of the cells, visualizing Cdc19^4D^-GFP foci, in cells deleted (*gpa2τ<*) or not for *GPA2*, during stationary phase (D6) or 180 min after release into fresh medium. The percentage (%) of cells with foci +/- SEM were quantified from three independent experiments in 480-800 cells each condition; contrast/brightness adjusted to best visualize foci. **(C)** Biomass accumulation (in arbitrary units, A.U.) of wild-type (WT) cells treated with NM-PP1 (blue curve) or for control DMSO (green curve) was measured during 24hrs recovery from 6 day (D6) stationary phase. Recordings start after inhibitor wash out and addition of fresh media (time 0). Error bars depict SEM from 3 independent experiments. Note that NM-PP1 exposure does not affect recovery of wild-type cells, excluding unspecific effects and problems washing-out NM-PP1. **(D)** and (**E**) Exponentially growing *pka-as* cells treated (PKA Inh) or not (Ctrl) with NM-PP1 were kept at 30°C (**D**) or heat-shocked for 30 min at 42°C (**E**). Biomass accumulation (in arbitrary units, A.U.) was then measured during 20hrs after inhibitor wash out and return to fresh growth media at 30°C (time 0). Error bars depict standard error of the mean from independent experiments (Heat shock: n=3 each condition, Expo: n=4 each condition). **(F)** Principal component (PC) analysis based on the metabolite intensities determined in the metabolomics dataset (Table 4). Different conditions are highlighted with distinct colors. **(G)** Schematic representation of the major steps of glycolysis, the TCA cycle and the trehalose/glycogen metabolism. PEP: phosphoenolpyruvate.

**Figure S6:**
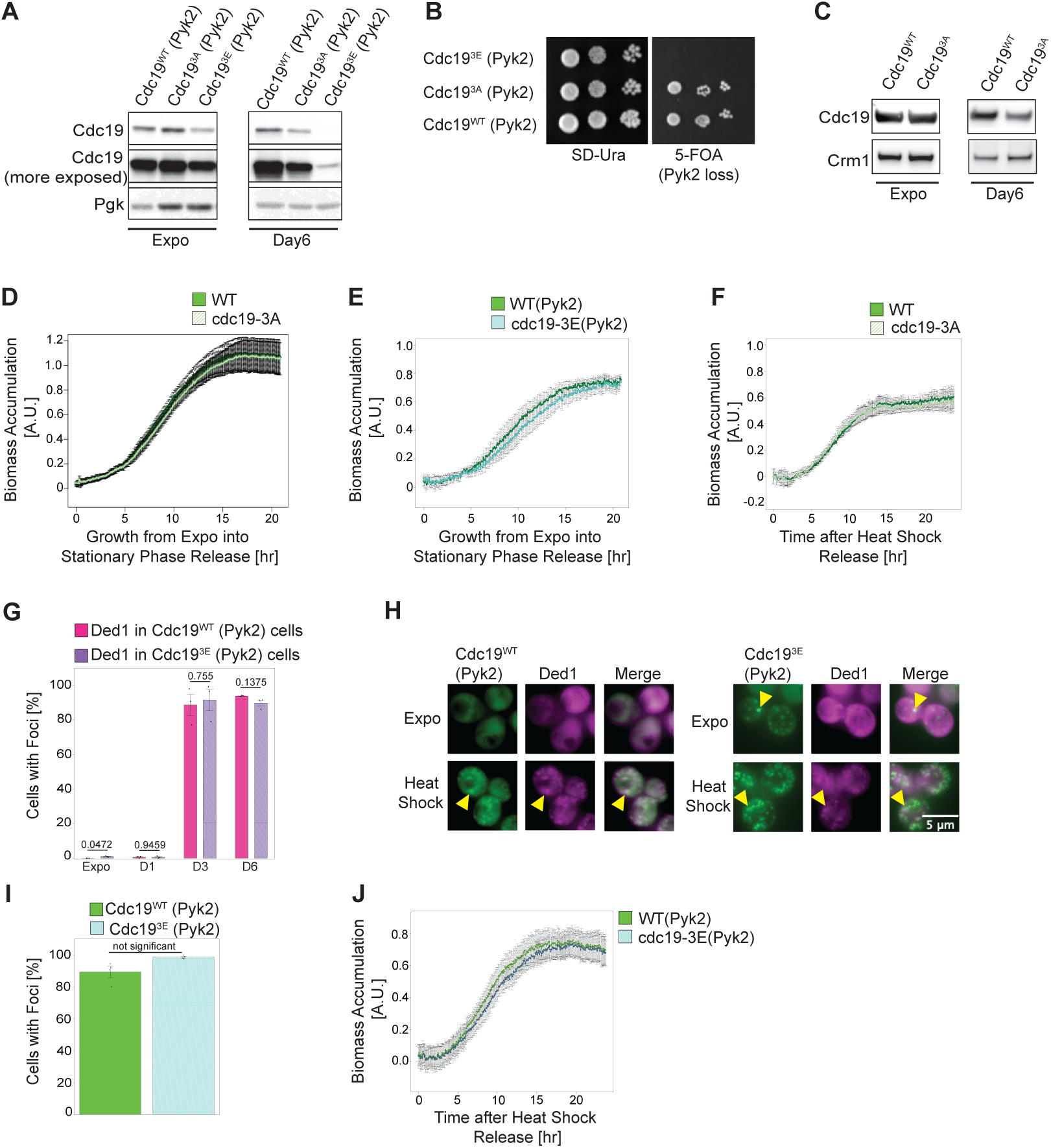
**(A)** Protein levels were compared by western blotting with GFP-antibodies of cell extracts prepared from cells expressing either GFP-tagged wild-type Cdc19 (Cdc19^WT^), non-phosphorylatable Cdc19^3A^ (S56A, S70A, S175A) or phosphomimicking Cdc19^3E^ (S56E, S70E, S175E). Cells were harvested from exponential growth (Expo) or 6-day stationary phase (Day6). All cells overexpress Pyk2 to ensure comparable growth rates. An antibody recognizing Pgk1 controls equal loading. A representative blot of 3 independent replicates is shown. **(B)** Serial dilution spottings of wild-type (WT), non-phosphorylatable *cdc19-3A* or phosphomimicking *cdc19-3E* mutant cells growing on nutrient-rich media with (left plate, SD-URA) or without (right plate, 5-FOA) a plasmid overexpressing Pyk2. A representative experiment from 3 independent replicates was photographed after 3 days at 30^0^C. Note that the phosphomimicking *cdc19-3E* mutant is unable to sustain growth in the absence of Pyk2. **(C)** Protein levels were compared by western blotting with GFP-antibodies of extracts prepared from cells without Pyk2 overexpression expressing either GFP-tagged wild-type Cdc19 (Cdc19^WT^) or non-phosphorylatable Cdc19^3A^ (S56A, S70A, S175A). Cells were harvested from exponential growth (Expo) or 6-day stationary phase (Day6). An antibody recognizing Crm1 controls equal loading. Representative blots of two (Day6) or three (Expo) replicates shown. **(D)** and (**E**) Biomass accumulation (OD=600) (in arbitrary units, A.U.) of cells expressing GFP-tagged wild-type (WT) Cdc19, non-phosphorylatable Cdc19-3A (S56A, S70A, S175A) or the phosphomimicking Cdc19-3E (S5E, S70E, S175E) mutant at different hours (hr) of growth from exponential phase (Expo) into stationary phase. Where indicated, the cells also overexpress Pyk2. Error bars depict standard error of the mean. Shown is the comparison of (**D**) n=6 replicates for Cdc19-WT and Cdc19-3A and (**E**) n=3 replicates for Cdc19-WT (Pyk2) and Cdc19-3E (Pyk2). **(F)** and **(J)** Biomass accumulation measurements (in arbitrary units, A.U.) of cells comparing either (**F**) GFP-tagged wild-type (WT) Cdc19 and the non-phosphorylatable Cdc19-3A or (**J**) wild-type (WT) Cdc19 and the phosphomimicking Cdc19-3E mutant after hours of recovery (hr) from heat stress of 30 min at 42^0^C. Where indicated, the cells also overexpress Pyk2. Error bars depict standard error of the mean from three replicates in independent experiments. **(G)** Wild-type or *cdc19-3E* cells overexpressing Pyk2 and harboring Ded1- mCherry (Ded1) were analyzed microscopically when exponentially growing (Expo) or arrested for different days into stationary phase (D1, D3, D6). The percentage (%) of cells with Ded1-mCherry foci was quantified, and standard error of the mean calculated from 3 independent experiments using 300-600 cells for each condition. Statistical significance was determined by an unpaired t-test. **(H)** and **(I)** Cells expressing Ded1-mCherry (Ded1) and harboring either GFP-tagged wild-type (Cdc19^WT^, left panel) or the phosphomimicking Cdc19^3E^ (S56E, S70E, S175E) mutant (right panel) were analyzed microscopically (**H**) when exponentially growing (Expo) or after heat stress for 30 min at 42^0^C (Heat Shock). Cell viability was maintained by co-expressing Pyk2. The arrowhead marks cytoplasmic foci. Representative images of 3 replicates are shown; contrast/brightness settings equal in same color channel and condition. Scale bar: 5µM **(I)** The percentage (%) of cells with GFP-tagged Cdc19^WT^ or Cdc19^3E^ foci were quantified after heat stress for 30 min at 42^0^C (Heat Shock). SEM were calculated from three independent experiments counting 530-560 cells for each condition. An unpaired t-test revealed no statistically significant difference (p=0.06) **(I)** See under Figure legend S6F.

**Figure S7:**
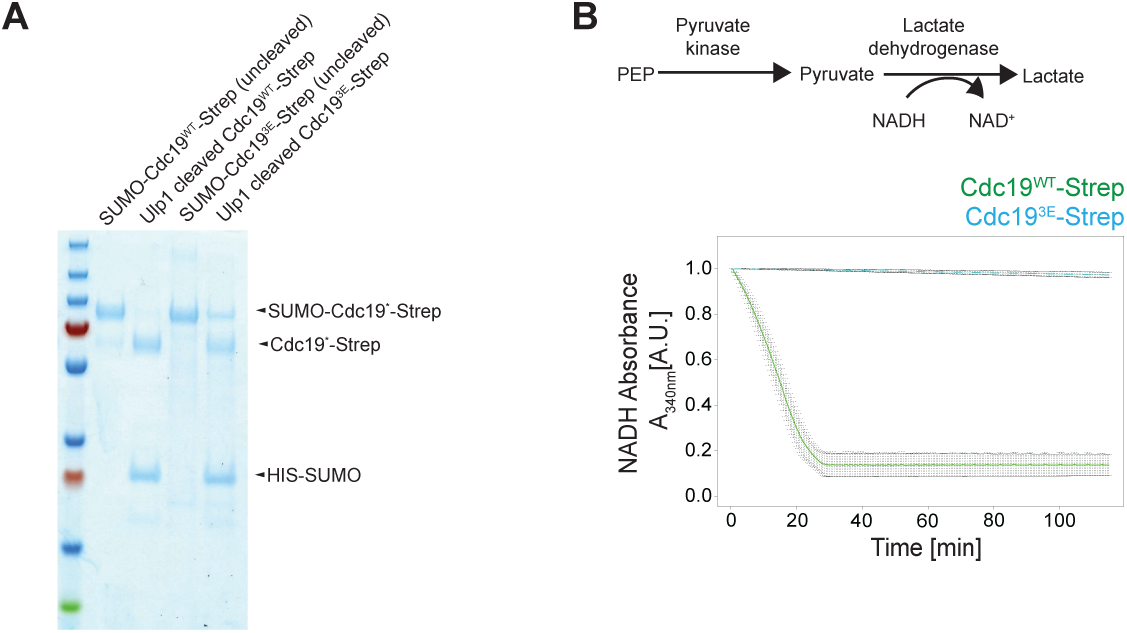
**(A)** Wild-type (WT) Cdc19 and the phosphomimicking Cdc19-3E mutant fused with a SUMO-Strep-tag were expressed in *E. coli*, purified over an affinity column, and analyzed by SDS-PAGE and Coomassie staining. The arrow points to the Cdc19 fusion proteins, which migrate at the expected size based on the protein markers. For the assays shown in Figures 7C-7G and S7B, the SUMO-tag was cleaved off by Ulp1 protease. A representative image of two replicates is shown. **(B)** The catalytic activity of Strep-tagged wild-type Cdc19 (WT) and the Cdc19-3E mutant protein after Ulp1 protease cleavage were measured in three replicates using a lactate dehydrogenase-coupled assay. Briefly, active Cdc19 converts phosphoenolpyruvate (PEP) into pyruvate, which in turn is reduced to lactate by lactate-dehydrogenase. The concomitant conversion of NADH to NAD^+^ is quantified as decrease in 340nm absorbance over time (minutes) and used as a measure of pyruvate kinase activity. Note that the Cdc19-3E mutant enzyme is catalytically inactive.

## Materials and Methods

### Yeast cell growth and fluorescence microscopy

Cells were grown in synthetic SD media (2 % glucose, 0.5 % NH4-sulfate, 0.17 % yeast nitrogen base (Sunrise Science Products, Cat# 1523-100), and amino acids) at 30 °C (see Suppl. Table S4 for all strains). To observe growth, cells were either (1) spotted in serial dilutions on SD agar plates and the plates imaged after 3 days at 30 °C, or (2) cells were washed once in growth media, loaded at OD600 0.1 into 24-well plates (Nunc) and optical density at 600 nm analyzed at different times using a Clariostar plate reader (BMG Labtech). Fluorescence microscopy was performed using a Nikon Eclipse Ti-E microscope with NIS-Elements Advanced Research V 5.02 software. For time-lapse experiments, yeast cells were loaded in CellASIC microfluidic chips (Merck Millipore, Cat# Y04C-02-5PK) and images recorded every 5 min. Budding index was calculated using the Cell Counter plugin (Fiji ^85^). To assess cell death, propidium iodide (Sigma, Cat# 81845-100MG) dye was used at 2.5 µg/ml dilution. The average cell size was measured using Fiji thresholded images, analyzing particles between 0.3-4.00 units.

### Quantitative phase imaging (QPI)-microscopy

For imaging stress granules during long-term stationary phase, we used refractive index tomograms with 3D fluorescence imaging capability (HT-X1; Tomocube Inc.). To reveal foci position, we scanned the sample in fluorescence (FL) mode to acquire images of the Cdc19-GFP signal (z-stack spacing=0.37 µm). For deconvolution of the raw data, we used the built-in Tomocube software (TomoStudio^TM^). To acquire refractive index (RI) images, we acquired 240 images with incoherent light illumination^86^ and reconstituted the RI tomograms using the built-in software (TomoStudio^TM^). The pixels of FL and RI images were re-sized to overlay each other to find the foci.

### Fluorescence recovery after photobleaching (FRAP) measurements

Images were acquired using a Yokogawa W1 Spinning disk on a Nikon eclipse Ti2 inverted microscope, using a Nikon 100x 1.45 CFI Plan Apo Oil objective. Acquired images had pixel sizes of 0.065 µm. The images were collected with an ORCA Fusion Digital CMOS camera (C14440-20UP, Hamamatsu), with an emission filter 525/50 and a 200mw 488nm diode laser (200mW). Images before and after photobleaching were acquired with 150 ms excitation time. The GFP signal was bleached in a 1.36 × 1.36 µm area using the 480 nm diode laser at 100% intensity. Image analysis was performed with the TrackMate Plugin from the Fiji image processing software with an estimated object diameter of 1µm. The signal from three areas was obtained for each time point: the bleached foci (IF), a reference foci in a neighboring cell (IR) and an area of equal size in the background (IB). The fluorescence signal in the foci was normalized as follows ^87^:

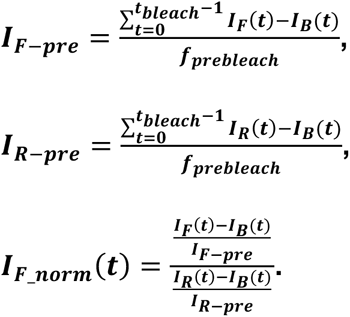

To reveal the half-time of recovery, we fitted the values of IF_norm with:

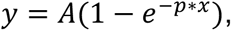

where *p* is the time constant and *A* is the fluorescence intensity.

The half time t1/2 was calculated using:

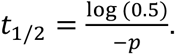

Data were analyzed, tested for statistical significance, and plotted using R software. Boxes in boxplots extend from the 25th to 75th percentiles, with a line at the median.

### Cloning

DNA mutations were introduced by site-directed mutagenesis using standard molecular biology protocols and controlled by sequencing.

### Semi-denaturing detergent agarose gel electrophoresis (SDD-AGE)

Yeast strains expressing GFP-tagged proteins of interest were grown for the indicated times in 5 ml media at 30 °C. After harvesting, cells were washed once with water and resuspended in ∼300 µl ice-cold lysis buffer (50 mM Tris pH 7.5, 150 mM NaCl, 1 % (vol/vol) TritonX-100, 2.5 mM EDTA, 0.33 mM PMSF, protease inhibitor tablet (Roche, Cat# 11836170001), 6.7 mM NEM). The mixture was added to ice-cold glass beads and the cells were lysed using mechanical disruption (6 m/s for three times 20 s with 5 min pause). After centrifugation, the supernatant samples were adjusted for equal protein concentrations and mixed 4:1 with 4 × Sample buffer (40 mM Tris acetic acid, 2 mM EDTA, 20 % glycerol, 4 % SDS (Cat# 436143-100G, Sigma), bromophenol blue). Samples were incubated for 10 min at room temperature and loaded onto a 1.5% agarose gel containing 0.1% SDS in 1 × TAE/0.1% SDS running buffer. The gel was run at low voltage in the cold. Proteins were detected by immunoblotting with a GFP-specific antibody.

### Size-exclusion chromatography (SEC)

For size exclusion analysis, yeast strains expressing GFP-tagged proteins of interest were grown in 50ml to OD=0.5. After harvesting, cells were washed once with water and resuspended in ice-cold lysis buffer (50 mM Tris pH 7.5, 150 mM NaCl, 2.5 mM EDTA, 0.5% Tween 20, 10% Glycerol, 1mM NaF, 1mM Na vandate, 20mM beta-Gly, 0.4mM PMSF, 8mM NEM, protease inhibitor). The mixture was added to ice-cold glass beads and the cells were lysed using mechanical disruption (6 m/s for three times 20 s with 5 min pause). After centrifugation, the supernatant was applied onto 0.22µm SpinX-Centrifuge Filters and about 2-4mg total protein was loaded on a Superdex 200 10/300 GL size-exclusion column (GE Healthcare) connected to an ÄKTA pure (GE Healthcare) at 4 °C. Protein elution was followed by measuring UV absorbance (280 a.u.) and fractions containing Cdc19 were analyzed by immunoblotting with GFP antibodies.

### Protein levels quantification & Sedimentation analysis

To prepare total protein extracts, yeast strains expressing GFP-tagged proteins of interest were harvested and lysed as described for SDD-AGE. After centrifugation, supernatants were analyzed by western blotting using α-GFP (Roche, Cat# 11814460001), α-RPS6-P (Cell Signaling, Cat# 4856), α-Pgk1 (Life technologies, Cat# 459250) or α-Crm1 (gift from Karsten Weis lab, ETH Zurich) antibodies. For sedimentation assays, aggregates were separated by centrifugation (10 min, 21’000g, 4°C) from the supernatant, and the input, pellet and supernatant fractions were analyzed by western blotting.

### Luminescence reaction to determine ATP levels

Metabolites were extracted as described under Metabolite-enrichment MS, solubilized in water and sonicated. Samples and CellTiter-Glo 2.0 assay reagents (Promega, Cat# G9241) were mixed 1:1, incubated for 10 min and the luminescence signal was recorded in 96-well plates (Greiner, F-bottom) in a Clariostar plate reader.

### Glucose Uptake Measurements

Glucose Content in the yeast media was measured using a glucose assay kit (Megazyme, Cat# K-GLUHK-110A) following the manufacturer’s protocol.

### Screen for suppressors of irreversible Cdc19 aggregates

Cells (BY4741) expressing the irreversible Cdc19-4D mutant (*cdc19-T372D, T376D, S377D, S385D* ^15^) were used in a SATAY transposon screen to identify genes that allow Cdc19-4D re-solubilization ^88^. Briefly, galactose-inducible transposase expression in *cdc19-4D* cells was induced for 15 days from a plasmid on plates containing 2% galactose. After harvesting, the cells were grown until saturation and heat-shocked (HS) or not at 42°C for 30 min. Genomic DNA was extracted, digested with Sau3AI, and inserted transposons identified by Ilumina sequencing. Transposons per gene and reads per gene in control versus HS conditions were compared, and significantly enriched (P<0.05) suppressors ranked by statistical significance.

### Phospho-enrichment mass-spectrometry Sample preparation

Cells were grown in 50 ml SD media until OD 5 for stationary phase and 500 ml of SD medium until OD 0.5 for logarithmic phase. Where indicated, 200 nM rapamycin (LC-Laboratories, Cat# R-5000) was added for 2 hrs. To analyze PKA-dependent phosphorylation sites, yeast *pka-as* cells were grown in 50 ml culture media to OD 5 (for Day 8 stat. phase) and 500 ml culture media to OD 0.5 (for Heat Shock and Expo. Phase), when indicated, cells were treated with 10 nM NM-PP1 (Cat# 13330, Cayman Chemical) to inhibit PKA. Cultures were quenched with 6.25% (v:v) of 100% trichloroacetic acid (TCA, Roth, Cat# 3744.1) on ice for 10 min, centrifuged at 1500g for 5 min at 4°C, pellets washed twice with ice-cold acetone, and stored at -80°C until lysis. Cell pellets were mechanically lysed at 4°C in 0.4 ml of 8 M Urea (Sigma, Cat# U5378), 100 mM ammonium bicarbonate (Sigma, Cat# A6141), 5 mM EDTA by glass bead beating (using beads with 500 µm diameter, equal volume), using the following settings: 6 m/s for 20 sec for 5 cycles with fresh lysis buffer, keeping the supernatant for every cycle. Protein concentrations were estimated by BCA Protein Assay (Thermo, Cat# A65453), and 3 mg of extracted proteins reduced with 5 mM TCEP (Cat# 51805-45-9, Sigma) for 30 min at 37°C, followed by 12 mM IAA (Indole-3-acetic acid, Sigma, Cat# I1149) at room temperature in the dark. After diluting Urea to 1 M by adding 100 mM Ammonium Bicarbonate, proteins were digested with 1:50 sequencing-grade Trypsin (Promega, Cat# V5111) overnight at 37°C and 300 rpm. Proteolysis was quenched by adding formic acid to a final volume of 1%, and the peptide mixture was cleaned by C18 phase separation cartridges (Waters, Cat# WAT020805). Briefly, peptides were loaded in the column, washed with methanol (1 column volume (CV)) and 50% acetonitrile (Sigma, Cat# 1000291000) /0.1% formic acid (1 CV), and equilibrated with 0.1% formic acid. After washing with 3 CV 0.1% formic acid, peptides were eluted with 2 CV 50% acetonitrile/0.1% formic acid, dried in a vacuum centrifuge, and resuspended in 50 µl 0.1% formic acid. 2 µl peptides were further diluted to 1 µg/µl and analyzed by LC-MS to normalize the phosphorylated level of non-enriched samples. For phospho-enrichment, the remaining peptides (48 µl) were diluted with 350 µl phospho-enrichment loading buffer (50% acetonitrile, 29.2% lactic acid and 0.1% trifluoracetic acid (TFA, Cat# T6508, Sigma)), and incubated in a tube rotator for 1 hour at room temperature with 6 mg of TiO2 resin (GL Science, Cat# 5010-21315), washed twice with 500 µl of methanol and twice with 500 µl of the loading buffer. The TiO2 resin supernatant was removed by centrifugation at 2000g for 1 min and the resin washed sequentially twice each with 500 µl of loading buffer, 500 µl of 50% acetonitrile, 0.1% TFA and finally 500 µl of 0.1% TFA. Phosphopeptides were eluted in two steps with 150 µl 50mM ammonium hydroxide (pH 10.8), quenched immediately with 4.4 µl 50% TFA, desalted using ultra-micro spin columns (The Nest Group, Cat# SUM S010-S100) and dried on a speedvac. Peptides were reconstituted to 1 µg/µl in 0.1% formic acid and retention time standard peptide (1:50 *iRT,* Biognosys, Cat# 1816351) were spiked before LC-MS analysis.

### LC-MS data acquisition

Phospho-enriched peptides were analyzed in an Orbitrap Fusion Lumos (Thermo) equipped with a nano electrospray ion source coupled to ACQUITY UPLC system M-Class (Waters). Samples were acquired in DDA mode for the library generation and in DIA mode for quantification of (phospho)-precursors. Peptides were separated at 50°C on a 75µm-diameter capillary column packed with ReproSil-Pur C18-AQ resin (Dr. Maish, Cat# r119.aq., 1.9 µm). Peptides were eluted at 300 nl/min over a linear gradient of 2 hours (5-35% of Buffer A (0.1% formic acid) and Buffer B (100% acetonitrile, 0.1% formic acid). The DDA data acquisition mode was set to perform one MS1 scan followed by a MS2 scans for a cycle time of 3 sec; MS1 scans (R=120’000 at 400 m/z, AGC=400’000 and maximum IT= 10ms), HCD fragmentation (NCE=28%), isolation windows (1.6 m/z) and MS2 scans (R=30’000 at 400 m/z, AGC=100’000 and maximum IT=54ms). A dynamic exclusion of 30 s was applied and charge states lower than two and higher than seven were rejected for the isolation.

Data-Independent Acquisition (DIA) measurements: samples were analyzed with the same LC set up used for assay library generation. The mass spectrometer was operated in DIA mode with the following parameters: one full FTMS scan (350-2’000 m/z) at 120’000 resolution, 100 ms injection time and 20’000 AGC target, followed by 41 variable windows from 350 to 1’150 m/z with 1 m/z overlap at 30’000 resolution, 54 ms injection time and 500’000 AGC target. Precursor ions were fragmented with HCD, Normalized Collision Energy 28%. Non-enriched peptides were analyzed in an Orbitrap QE Executive Plus equipped with a nano electrospray ion source coupled to EASY-nLC-1000 liquid chromatography system (Thermo). Peptides were separated on a reverse phase column (75 µm ID x 400 mm New Objective, in-house packed with ReproSil Gold 120 C18, 1.9 µm) across a 120 min linear gradient from 5 to 30% (buffer A: 0.1% (v/v) formic acid; buffer B: 0.1% (v/v) formic acid, 95% (v/v) acetonitrile). The DDA data acquisition mode was set to perform one MS1 scan followed by a 30 MS2 scans for top 20 ion peaks; MS1 scans (R=70’000 at 400 m/z, AGC=100’000 and maximum IT= 55ms), HCD fragmentation (NCE=30%), isolation windows (1.4 m/z) and MS2 scans (R=17’500 at 400 m/z, AGC=100’000 and maximum IT=55ms). A dynamic exclusion of 30 s was applied and charge states lower than two and higher than seven were rejected for the isolation.

Targeted MS analysis: the reference phosphopeptides (PEPotec Heavy, Thermo) for Atg1 S515, Bcy1 S145, Cdc19 S22, Cdc19 S56, Cdc19 S9, Cdc19 S407, Msn2 S288 and iRT QC peptides with arginine [+10] or lysine [+8] labeled with heavy amino acids were synthesized and spiked into the sample mixture. Samples were acquired with same instrument setup in untargeted (DDA) mode and in targeted mode for quantification using a shorter gradient (peptides were eluted in 60 min from 5 to 40% buffer B). A targeted scheduled method was set up with one MS1 scan (R=70’000 at 400 m/z, normalized AGC = 3’000’000 and max. IT 100 ms) and a cycle time of 3s to monitor 69 peptides (R= 35’000, AGC = 50’000 and max. IT 110 ms) using an isolation window of 1.8 m/z and normalized collision energy 27%.

### Analysis of MS data

Spectra libraries for precursors and phosphoprecursors were generated based on 20 DDA runs pooling the samples generated for DIA acquisition (pooled conditions: exponential growth, days 1-5, day 6, days 7-8, rapamycin). For each condition phospho-enriched and not enriched pooled samples were acquired in two technical replicates. The spectra from mzXML files (converted with msconvert) were searched using Comet search engine (version 2018.01) against a fasta file generated from SGD database (downloaded on 13.01.2015, 6’713 proteins) concatenated with decoy protein entries. The search parameters were as follows: ± 25 ppm peptide mass tolerance, semi-tryptic specificity, two missed cleavages allowed, cysteine carbamidomethylation as fixed modification and methionine oxidation as variable modifications. For the phospho-enriched analysis, phosphorylation of serine, threonine, and tyrosine residues were considered as variable modification. Results were further processed using peptideProphet and filtered at 1% FDR (PSM level). A consensus library for phospho-enriched (ConsPhosphoLib_luciphor_openswath_curated_concat.tsv) and not enriched (ConsNonEnrichedLib_openswath_curated.tsv) datasets were generated using SpectraST using the following parameters: 5 most intense fragments per peptide (charge 1+ or 2+), y or b fragment ions for each spectrum within the mass range 350-2000 m/z. The generated library for the phospho dataset contained 57’870 precursors of which 46’372 phosphoprecursor, while the library for non-phospho enriched sample comprised 45’556 precursors for a total of 4’488 protein groups. In the following step, DIA data were extracted with Spectronaut (v. 12.0.20491.11.25470 ^89^) using the default settings, except for the following options: machine learning defined as “across experiment”; no “cross run normalization” and protein inference “From protein-db matching”. Spectronaut quantification is reported for phospho and not enriched dataset at precursors and fragments ion levels (“*20190405_163545_ Phospho_Report.xls*” and “*20190522_161058_ NE_Report.xls”*, respectively for phospho and non-phospho dataset). For the analysis of phosphorylated peptides, precursor intensities identified in at least two replicates of one condition were log2 transformed and median normalized, excluding precursors without phosphorylation modifications. Missing values were imputed per precursor by sampling from a normal distribution centered at 𝑥=𝑥-1.8 × 𝑠𝑑, where 𝑥 is the mean value of the precursor and sd is the standard deviation across all conditions for each precursor. The analysis of variance (ANOVA) for each phosphoprecursor intensity and condition compared to the mean across all conditions was applied to filter phosphoprecursors that change significantly in at least one condition. Phosphoprecursors showing significant changes in at least one condition over the mean of all conditions (|log2FC| > 1 and p-value adjusted with Benjamin Hochberg correction < 0.01) were considered in the analysis (only 26’450 were considered, out of 43’828 phosphoprecursors). Phosphopeptide abundance (22’121) was calculated from the average intensity of filtered phosphoprecursors. The average of phosphopeptides per condition was normalized for the average abundance per condition of proteins containing the phosphoprecursor. After protein normalization, 16’908 phosphopeptides from 1’564 proteins were considered for unsupervised hierarchical soft cluster (fuzzy c-means clustering) using the R package Mfuzz ^90^. The number of clusters (12) was determined by minimizing intra-cluster variance and maximizing the average silhouette. Before clustering, data were standardized to ensure that all normalized phosphopeptides across all conditions had a mean of 0 and standard deviation of 1. Only phosphopeptides with a minimum membership threshold of 0.6 were assigned to a specific cluster, otherwise they were assigned as “cluster NA”. To calculate protein abundance from the non-enriched dataset, intensities of precursors were log2 transformed and then normalized using the median across all runs. Precursors identified sparsely across the matrix were filtered out (44’072 precursors identified in duplicate in at least one condition being considered). Missing values were imputed as described above for the phosphorylation dataset. Protein abundance for each condition was determined by averaging the levels of all identified precursors per protein, considering only proteins with at least three precursors. Overall, protein levels were inferred for a total of 2’711 proteins. Each cluster was further characterized by gene ontology enrichment analysis ^91^, performing a hypergeometric test with the background set consisting of the clustered phosphopeptides. Enrichment significance for stress granule components was assessed using a protein list identified as more abundant in the Pab1 BioID experiment under starvation conditions compared to control, as described in Amen *et al*. ^41^. Additionally, enrichment significance for phosphosites localized in intrinsically disordered regions was calculated by identifying IDR regions in the yeast proteome, defined by the consecutive presence of at least five amino acids with an IUPRED score greater than 0.8 ^42^. Kinase-substrate relationships for all clusters were inferred using the NetworKIN 3.0 algorithm ^43^, considering relationships with a score higher than 4. Subsequently, kinase-substrate enrichment in each cluster was determined using a hypergeometric test, using as background the clustered phosphosites. Motif analysis for cluster nine was further confirmed by the MotifAll algorithm (PhosphoSitePlus.org), using a list of 15-mers centered on the phosphorylated site. The same list of 15-mers with scrambled sequences was used for control. The entire dataset, including raw data, generated tables and scripts used for the data analysis is available in the PRIDE repository (PXD053013).

The analysis of MS data of PKA-dependent sites was performed as follows: acquired DDA spectra in the untargeted analysis were searched using the MaxQuant software package embedded with the Andromeda search engine ^92^ against the *S. cerevisiae* reference dataset (http://www.uniprot.org/, downloaded on 17.05.2021, 6055 entries) extended with reverse decoy sequences. The search parameters were set to include semi-tryptic peptides, maximum two missed cleavage, carbamidomethyl as static peptide modification, oxidation (M), deamidation N terminal, phosphorylation (S, T, Y) as variable modification and “match between runs” option. The MS and MS/MS mass tolerance were set to 20 ppm and 0.5 Da, respectively. False discovery rate of < 1% was used at the PSM level. Phosphorylation sites identified in the table “Phospho (STY)Sites.txt” were filter for localization (localization score > 0.6) and only phosphosites identified at least in two replicates in at least one conditions were considered (7’152 phosphosites identified). Intensity values of phosphorylation sites were log2 transformed, median normalized and imputed using random sampling from a normal distribution generated from the 1% lower intense values of the whole dataset. ANOVA statistical analysis was performed to identify significantly regulated phosphorylation sites (p-values were corrected for multiple hypothesis using Benjamin Hochberg method). Downregulated phosphosites (log2FC < (-1) & *p* value BH corrected <0.05) were characterized by gene ontology enrichment analysis (EnrichR package) with the identified proteins in the dataset serving as the background and calculating the significance of the enrichment of GO terms (downloaded from Quick GO database, 11.2024). The identification of kinase-substrate relationships was calculated by NetworKIN algorithm and the identification of kinases significantly enriched was assessed by hypergeometric test with the background set consisting of the identified phosphosites in the dataset. For targeted PRM analysis, peptides were manually examined using Skyline daily software ^93^. Endogenous phosphorylated peptides were identified by matching the coelution and peak shapes of precursor and product ions with those of the reference peptides. Quantification was carried out using data from the file “Transition_Results_240608.csv,” summing five fragment ions per peptide. The intensity of endogenous peptides was normalized to the intensity of the heavy peptides. After log2 transformation and normalization against the mean phosphorylation intensities in the exponential condition without treatment, the significance of changes was evaluated using p-values obtained from two-sided, unpaired t-tests. The entire dataset, including raw data, generated tables and scripts used for the data analysis is available in the PRIDE repository (PXD052971).

### *In vitro* kinase assay and MS analysis

*In vitro* kinase reactions were performed using recombinant Cdc19, which was mixed at a concentration of 0.5 mg/ml 1:50 with PKA (NEB, Cat# P6000S) in buffer containing 100 mM Tris-HCl pH7.5, 100 mM NaCl, 5% glycerol, 20 mM MgCl2, 3 mM DTT, 400 µM ATP (Sigma, Cat# A2383-5G), 1 mM PMSF with protease inhibitor tablets. The incubation was done for 1hr or 24 hrs at 30°C. For visualization the samples were loaded onto a SuperSep Phos-tag gel (Wako Chemicals, Cat# 192-18001) and stained by Coomassie. For MS analysis, reactions were quenched, ∼1 µg of purified Cdc19 was subjected to proteolysis and MS identification and quantification. Cdc19 was loaded on a Filter (Vivacon 500, 10MKCO, Sartorius, Cat# VN01H02) and centrifuged until dry (centrifugation 8000g, 15 min). After denaturation (8 M urea), reduction (5 mM tris(2-carboxyethyl)phosphine (TCEP)), and alkylation (10 mM iodoacetamide, Sigma, Cat# I5161), Cdc19 was washed with 25 mM ammonium bicarbonate (three centrifugation steps, 8000g, 15 min) and overnight proteolyzed (0.2 μg of trypsin). Proteolysis was quenched by 0.1% formic acid, and dried peptides were resuspended in 10 μl of 0.1% formic acid and 2% acetonitrile supplemented with iRT peptides for quality control. Identification of Cdc19 phosphopeptides were performed by LC-MS/MS operating in data dependent acquisition (DDA) or parallel reaction monitoring (PRM) to identify Cdc19 S70 phosphopeptide (S[Pho]EELYPGRPLAIALDTK). For this peptide, fragments, transitions ions and relative elution time were extracted from the DDA library generated for the profiling of phosphorylation levels during starvation and rapamycin conditions (ConsPhosphoLib_luciphor_openswath_curated_concat.tsv).

Samples were acquired in untargeted (DDA) mode and in targeted mode to quantify Cdc19 and iRT QC peptides. For DDA and PRM analysis, samples were acquired in an Orbitrap Exploris 480 equipped with a nanoelectrospray ion source coupled to an Vanquish Neo liquid chromatography system (Thermo). Peptides were separated using a reverse phase column (75 µm ID x 400 mm New Objective, in-house packed with ReproSil Gold 120 C18, 1.9 µm) in a gradient of 60 min from 7 to 35% (buffer A: 0.1% (v/v) formic acid; buffer B: 0.1% (v/v) formic acid, 80% (v/v) acetonitrile). The DDA data acquisition mode was set to perform one MS1 scan followed by a 20 MS2 scans for top 20 ion peaks; MS1 scans (R=60’000 at 400 m/z, AGC targeted =100% and maximum IT= Auto), HCD fragmentation (NCE=28%), isolation windows (1.4m/z) and MS2 scans (R=30’000 at 400 m/z, AGC normalized=300% and maximum IT=Auto). A dynamic exclusion of 25s was applied and charge states lower than two and higher than seven were rejected for the isolation. The targeted scheduled method was set up with one MS1 scan (R=120’000 at 400 m/z, normalized AGC = 100% and max. IT = 256ms) and a cycle time of 3s to monitor 15 peptides (R= 256’000, AGC normalized = 200% and max. IT 110ms) using an isolation window of 1.8m/z and normalized collision energy 30%.

The DDA MS data were searched using the MaxQuant software package embedded with the Andromeda search engine ^92^ against the sequences of Cdc19, Tpk1, Tpk2 and Tpk3 (http://www.uniprot.org/, downloaded on 10.2022) extended with reverse decoy sequences. The search parameters were set to include semispecific tryptic peptides, maximum two missed cleavage, carbamidomethyl as static peptide modification, oxidation (M), deamidation N terminal, phosphorylation (S, T, Y) as variable modification and “match between runs” option. The MS and MS/MS mass tolerance were set to 20 ppm and 0.5 Da, respectively. False discovery rate of < 1% was used at the PSM level. Cdc19 phosphorylation sites listed in the “Phospho (STY)Sites.txt” table were filtered based on localization criteria (localization score > 0.8) and identification in all three replicates under the condition where Cdc19 was incubated with PKA for 24 hours. For targeted PRM analysis, S[Pho]EELYPGRPLAIALDTK (Cdc19 S70) intensity was manually examined using Skyline daily software ^93^. The identification was based on co-elution of precursors and at least 10 transition ions. Quantification was carried out using data from the file “Transition_Results_230915.csv,” summing the top five most intense fragment ions. The entire dataset, including raw data, generated tables and scripts used for the data analysis are available in the PRIDE repository (PXD052995).

### Metabolite-enrichment MS and analysis

Metabolite measurements were carried out in triplicates. Cells were grown as described above, and harvested after 6 days stationary phase, or 4, 8, or 12 hrs after re-feeding. Cells were harvested on a 0.45µm pore size PVDF filter (Sigma, Cat# P1813-100EA), which was then immediately frozen in a -20°C cold extraction solution (40:40:20 acetonitrile/ methanol/ water). After 2 hrs, the extraction solution was centrifuged (15 min, 4.000 rpm, 0°C) and the supernatant containing the metabolites was transferred to fresh tubes. Samples were dried in a Speed-Vac and stored at - 80°C. Prior to MS analysis, samples were resuspended in 90% ACN (Acetonitrile), and 5µl measured in a Q Executive mass spectrometer (Thermo) coupled to a Vanquish Horizon Binary Pump (Thermo) equipped with Premier BEH Amide column (Waters, 150mm x 2.1mm), applying a gradient of 10mM ammonium bicarbonate in water (A) and 10mM ammonium bicarbonate in 95% acetonitrile (B) from 99% B to 30% B over 12min. The metabolomics data set was evaluated in an untargeted fashion with Compound Discoverer software (Thermo). mzCloud and mzVault have been used to score fragmentation patterns and assign MS2-based identities to the features. Feature filtering parameters used were the following: Signal/noise > 3, mzCloud or mzVault match >50, ppm mass error within +/- 5ppm., match with in-house developed MS1_RT library within +/- 10sec., chromatographic peak and MS2 spectra quality.

### Protein purification

*E. coli* Rosetta pLys cells (Novagen, Cat# 71403) were transformed with plasmids expressing wild-type or mutant Cdc19. Cells were grown at 37 °C in TB media (23.6g/L yeast extract, 11.8 g/L tryptone, 9.4 g/L K2HPO4, 2.2 g/L KH2PO4, 0.4% Glycerol) containing antibiotics, and protein expression was induced at OD600 0.6 with 0.1 mM IPTG (Applichem, Cat# A1008,0050). After growth at 16-18 °C for 12 h, cells were harvested by centrifugation and the pellet resuspended in purification buffer (100 mM Tris-HCl pH 7.4, 200 mM NaCl, 1 mM MgCl2, 10 % glycerol, 1 mM phenylmethylsulfonyl fluoride (PMSF), 1 mM DTT), supplemented with protease inhibitor tablets and 75 U/ml of Pierce universal nuclease (Thermo, Cat# 88700), 25 U/m Benzonase (Sigma, Cat# 71205-3) and 33.33 µg/ml RNAse (Thermo, Cat# AM2294). Cell lysis was achieved by freezer milling (six cycles of 2 min cooling and 3 min grinding at setting 15 CPS), and the lysates cleared by centrifugation (4 °C, 30 min, 48000 g). The supernatant was loaded at 4 °C onto a Strep-Tactin Superflow Plus column (Qiagen, Cat# 30060) following the manufacturer’s instructions, proteins eluted with purification buffer containing 2.5 mM D-desthiobiotin (Sigma, Cat# D1411-1G), and analyzed by SDS-PAGE and Coomassie blue staining. Pure proteins were aliquoted and stored at -80 °C.

### Pelleting Assay and Atomic force microscopy (AFM)

Purified proteins were thawed on ice and diluted to a final protein concentration of 0.2 mg/ml in 100 mM Tris/HCl, 200 mM NaCl, 1 mM MgCl2, 10% glycerol, 1 mM DTT, 1 mM PMSF, pH 7.4. To the protein, SUMO protease (Sigma, Cat# SAE0067-2500UN) was added 1:50, diluted 1:1 in citric phosphate buffer pH7.4 (no Stress) or citric phosphate buffer pH 5.2 (low pH) and incubated for 1hr at 30°C. For pelleting assays, aggregates were separated at 10 min, 21’000 g, 4 °C form the soluble protein, and input, supernatant and pellet fractions were loaded on a SDS-PAGE gel and stained by Coomassie. For AFM analysis, an additional low pH sample was heat shocked at 42°C for 1hr (low pH + HS). The freshly cleaved mica was functionalized by 1% (3-Aminopropyl)triethoxysilan (APTES, Sigma, Cat# 440140-100ML) for 2 min, rinsed with Milli-Q water and dried by compressed gas. An aliquot (10 µl) of diluted sample was deposited on the APTES-functionalized mica for 2 min, rinsed with Milli-Q water and dried by a gentle flow of compressed gas. AFM measurements were carried out using a multimode 8 AFM (Bruker) with an acoustic hood to minimize vibrational noise. AFM imaging was operated in soft tapping mode under the ambient condition, using a commercial silicon nitride cantilever at a vibration frequency of 70 kHz. AFM images were flattened using Nanoscope 8.1 software (Bruker), and further statistical analysis was carried out on the Fiberapp software ^94^.

### ThioflavinT (ThT) staining

Purified Cdc19^WT^ or mutant Cdc19^3E^ in purification buffer was thawed on ice, cleared by centrifugation (10 min, 4 °C, 21000 g), and the concentration adjusted to 0.15 mg/ml. Filtered (0.2 µm filter) Thioflavin T (Sigma, Cat# T-3516-25G) solution was added at 1:10 dilution in a 384-well plate (Sigma, Cat# CLS3766-100EA), and ThT binding measured in a CLARIOstar plate reader, with 450 nm excitation and 490 nm emission.

### Pyruvate kinase activity assay

The activity of purified Cdc19^WT^ and Cdc19^3E^ were measured as described ^24^. Briefly, the pyruvate kinase reaction was coupled to the lactate dehydrogenase reaction and the conversion of NADH to NAD^+^ measured at 340 nm. Purified proteins were diluted in activity buffer (50 mM imidazole pH 7, 100 mM KCl, 25 mM MgCl2, 10 mM ADP, 0.3 mM NADH, 10 U/ml LDH) to a final protein concentration of 2 μg/ml. Reactions were started by adding PEP (final concentration 2 mM), and the time-dependent decrease at 340 nm absorbance was monitored.

### Quantification and statistical analysis

All data shown in this paper are representative results from at least three independent experiments, unless specified differently in the figure legends. R was used to analyse and plot the results. Statistical details of experiments, number of independent replicates (n), *p*-values and dispersion and precision measures (mean and SEM) are indicated in the respective figure legends. T-tests were calculated as an unpaired test with unequal variance.

